# Giant extrachromosomal element “Inocle” potentially expands the adaptive capacity of the human oral microbiome

**DOI:** 10.1101/2024.09.30.615745

**Authors:** Yuya Kiguchi, Nagisa Hamamoto, Yukie Kashima, Lucky R. Runtuwene, Aya Ishizaka, Yuta Kuze, Tomohiro Enokida, Nobukazu Tanaka, Makoto Tahara, Shun-Ichiro Kageyama, Takao Fujisawa, Riu Yamashita, Akinori Kanai, Josef S. B. Tuda, Taketoshi Mizutani, Yutaka Suzuki

**Affiliations:** Department of Computational Biology and Medical Sciences, Graduate School of Frontier Sciences, The University of Tokyo, Tokyo, Japan; Laboratory for Symbiotic Microbiome Sciences, RIKEN Center for Integrative Medical Sciences, Yokohama, Japan; Medical & Biological Laboratories Co., Ltd., Tokyo, Japan; Life Science Data Research Center, Graduate School of Frontier Sciences, The University of Tokyo, Tokyo, Japan; AIDS Research Center, National Institute of Infectious Diseases, Tokyo, Japan; Division of Infectious Diseases, Advanced Clinical Research Center, the Institute of Medical Science, The University of Tokyo, Tokyo, Japan; Department of Head and Neck Medical Oncology, National Cancer Center Hospital East, Kashiwa, Japan; Division of Cancer Immunology, Research Institute/ Exploratory Oncology Research and Clinical Trial Center, National Cancer Center, Kashiwa, Japan; Department of Radiation Oncology, National Cancer Center Hospital East, Kashiwa, Japan; Division of Radiation Oncology and Particle Therapy, National Cancer Center Hospital East, Kashiwa, Japan; Translational Research Support Office, National Cancer Center Hospital East, Kashiwa, Japan; Division of Translational Informatics, Exploratory Oncology Research and Clinical Trial Center, National Cancer Center, Kashiwa, Japan; Faculty of Medicine, Sam Ratulangi University, Kampus Unsrat, Bahu Manado, Indonesia; Center for Emergency Preparedness and Response, National Institute of Infectious Diseases, Tokyo, Japan

## Abstract

Survival strategy of bacteria is expanded by extrachromosomal elements (ECEs). However, their genetic diversity and functional roles for adaptability are largely unknown. Here, we discovered a novel family of intracellular ECEs using 56 saliva samples by developing an efficient microbial DNA extraction method coupled with long-read metagenomics assembly. Even though this ECE family was not hitherto unidentified, our global prevalence analysis using 476 salivary metagenomic datasets elucidated that these ECEs reside in 74% of the population. These ECEs, which we named, “Inocles”, are giant plasmid-like circular genomic elements of 395 kb in length, having *Streptococcus* as a host bacterium. Inocles encode a series of genes that contribute to intracellular stress tolerance, such as oxidative stress and DNA damage, and cell wall biosynthesis and modification involved in the interactions with oral epithelial cells. Moreover, Inocles exhibited significant positive correlations with immune cells and proteins responding to microbial infection in peripheral blood. Intriguingly, we examined and found their marked reductions among 68 patients of head and neck cancers and colorectal cancers, suggesting its potential usage for a novel biomarker of gastrointestinal cancers. Our results suggest that Inocles potentially boost the adaptive capacity of host bacteria against various stressors in the oral environment.

## INTRODUCTION

Bacteria residing in humans are exposed to various stressors, including nutrient competition, drugs for humans, antibiotics, and human immune responses. To survive these multiple stressors, bacteria expand their adaptive capacity by acquiring accessory genes corresponding to environmental stress tolerance. Mobile genetic elements (MGEs) are the major mechanism by which bacteria obtain accessory genes from other bacteria. Integrative and conjugative elements (ICEs) and transposons are intrachromosomal MGEs that transfer the accessory genes to host bacteria by integrating their genome into the host bacterial chromosome^1,2^. In addition, bacteria obtain accessory genes via extrachromosomal elements (ECEs), which are genetic elements independent of chromosomes^3^. Microbiological research targeting pathogenic bacteria has revealed that ECEs such as plasmid confer environmental stress resistance on host bacteria, including antibiotic resistance^2^. This infers that ECEs contribute to expanding the adaptive capacity of human commensal bacteria.

Several advanced strategies for predicting ECEs from metagenomic sequences have revealed that human commensal bacteria harbor diverse ECEs, such as bacteriophages (phages)^4–9^, plasmids^10^, and viroid-like elements^11^. However, current metagenomic analysis based on a limited set of reference genomes may overlook ECEs that could not be classified using existing knowledge and it is largely unknown the genetic diversity and functional roles of ECEs in human commensal bacteria.

Here, we identified the novel giant ECEs from the oral cavity of most individuals in the world, which are considered plasmid-like elements encoding multiple genes related to adaptation to intracellular and extracellular stresses. We have characterized this novel plasmid-like element, termed “Inocle,” focusing on its genetic and ecological significance. Moreover, we elucidated the significant associations of Inocles with human immune statuses and certain cancer types. This study demonstrates that human commensal bacteria adapt to changes in their habitat conditions and human physiological alterations utilizing giant ECEs.

## RESULTS

### Long-read metagenomic sequencing optimized for human saliva samples

Long-read sequencing improves metagenomic assembly for the reconstruction of ECEs with high completeness^12–16^; however, human saliva contains 85–95% human genomic DNA, making it difficult to obtain sufficient sequencing depth from microbes^17^. Therefore, we developed a simple method to extract high-molecular-weight DNA with reduced human genomic DNA from saliva samples. After collecting bacterial pellets by centrifugation, we removed extracellular DNA, which should be mostly human DNA^17^, by treating the bacterial pellet with nuclease under optimal conditions for nuclease activity. We refer to this method as the “preNuc”. After the treatment of preNuc, we extracted high-molecular-weight DNA using enzymatic bacterial cell lysis (**Figure 1A**). Notably, preNuc reduced the average percentage of human DNA from 90% to 40% in short-read sequences prepared from fresh and frozen saliva samples (**Figure 1B and Table S1**). The preNuc treatment resulted in a two-fold increase in the total microbial bases compared to that in preNuc-untreated samples when using the PromethION sequencer of Oxford Nanopore Technologies (**Figure 1C**) without shortened read length (**Figure 1D**). Moreover, Spearman’s *rho* for bacterial composition averaged 0.83 ± 0.08 at the species level between preNuc-treated and -untreated saliva samples (**Figure 1E**), indicating that our method easily reduces the human DNA in saliva samples and extracts high-molecular-weight DNA while minimally affecting bacterial composition.

**Figure 1.**
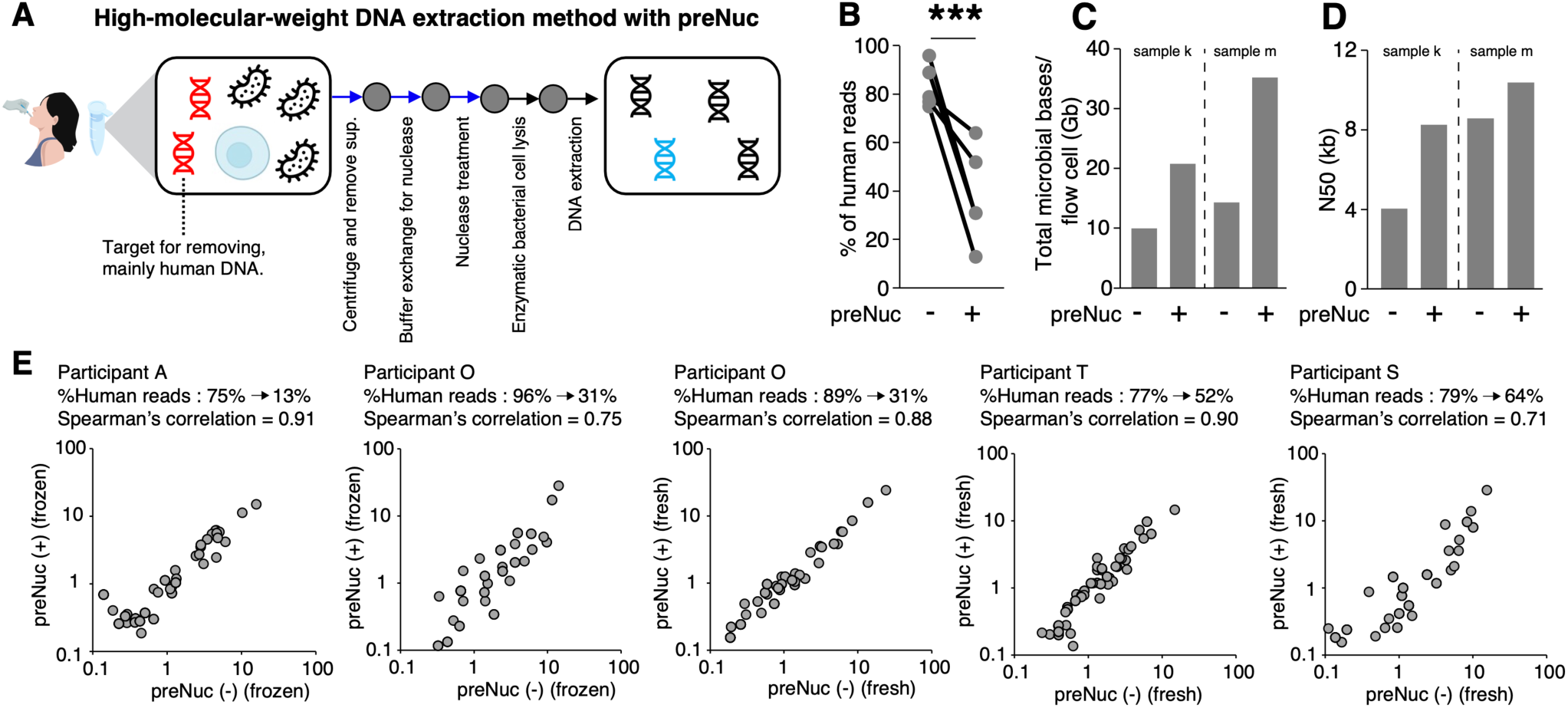
Impact of the preNuc treatment for salivary metagenomics. (A) Schematic workflow for high-molecular DNA extraction from human saliva samples. Blue arrows indicate processes of the preNuc. (B) The percentage of human reads in NovaSeq reads obtained using preNuc-treated and -untreated saliva samples. \*\*\**p* < 0.001, N.S., not significant based on paired student’s *t*-test. (C) Total bases of PromethION reads without low-quality and human reads per flow cell acquired from preNuc-treated and -untreated samples. (D) N50 length of PromethION reads without low-quality and human reads per flow cell of the preNuc-treated and -untreated samples. (E) The correlation of species-level bacterial composition between preNuc-treated and -untreated saliva samples.

Subsequently, we obtained PromethION sequences from 46 Japanese saliva samples with an average of 3.2 million high-quality and non-human reads, including 23.4 Gb with an N50 length of 12.5 kb per sample (**Table S2**). We performed *De novo* metagenomic assembly using metaFlye^18^ (**Table S3**) and used contigs with a depth exceeding 10 as high-quality contigs in subsequent analyses because they had an average of 98.95% similarity with the corresponding short-read contigs assembled using NovaSeq sequences (**Figure S1**).

### Discovery of unrecognized ECEs

We attempted to identify hitherto unrecognized ECEs to expand our knowledge of the genetic diversity of ECEs in the human oral microbiome. First, we classified 69,881 high-quality contigs, which are >3kb in length and unaligned to the human genome, into known genetic elements (**Figure S2A**) and classified 85.8% potential bacterial chromosomal, 1.5% phage, 2.9% plasmid, and 9.8% unclassified contigs (**Figure 2A**).

**Figure 2.**
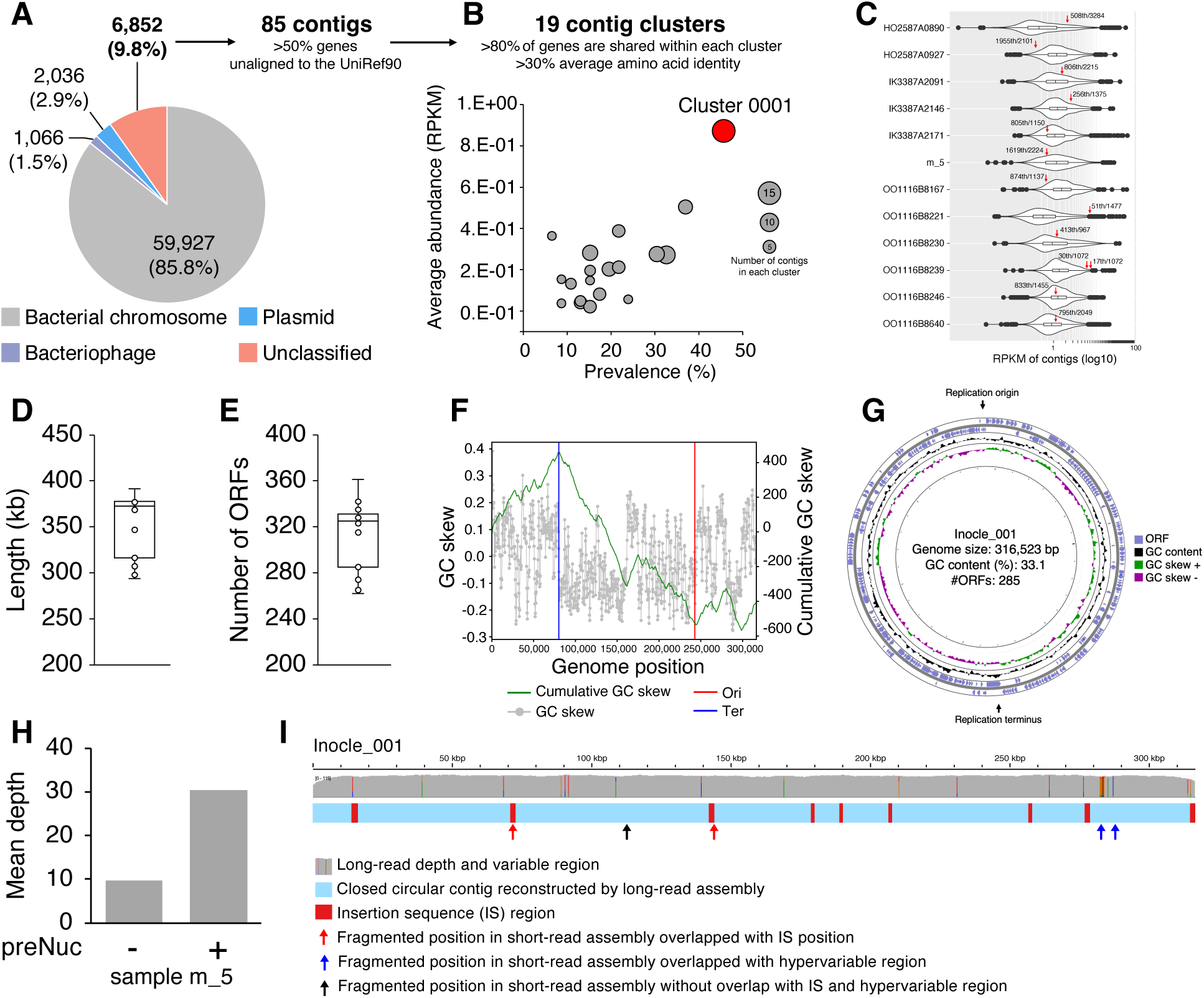
Discovery of potentially unrecognized genetic elements. (A) The ratio of each microbial genetic element of long-read high-quality contigs. Among the 6,852 unclassified contigs, 85 were further analyzed as strong candidates of unrecognized elements. (B) Average abundance (y-axis), prevalence (x-axis), and number of contigs in each operational contig cluster (plot size) obtained from 85 contigs. (C) Abundance of Cluster 0001 contigs among all contigs in each sample. The violin plot shows the distribution of the abundance of all contigs. The red arrow indicates the abundance of Cluster 0001 contigs and their rank in all contig abundance. Box plots represent the inter-quartile range (IQR), and lines inside the box indicate the median. Whiskers show 1.5 IQR. (D) The contig length and (E) number of ORFs of Cluster 0001. (F) Representative replichore structure of Cluster 0001 contig (Inocle_001). The green line indicates cumulative GC skew, and the gray plots indicate GC skew in each genomic position. The candidate replication origin and terminus are represented by vertical red and blue lines, respectively. (G) Representative genomic structure of Inocle: from inner to outside, GC skew, GC content, forward strand of ORFs, and reverse strand of ORFs. (H) The mean depth of the Cluster 0001 contig (Inocle_004) obtained from preNuc-treated and - untreated saliva samples. (I) Comparison of the long-read contig and corresponding short-read contigs. The blue bar represents the contig constructed by the long-read assembly, and the red boxes indicate IS regions. The arrows represent fragmented positions in short-read assembly. The upper gray bar plot indicates the depth of the mapped long reads. Colored bars in the depth bar plots indicate single-nucleotide variants: Green=A, red=T, orange=G, and blue=C.

Among 6,852 unclassified contigs, 85 were particularly considered unrecognized genetic elements with many unknown functional genes (**Figure 2A**). Specifically, over 50% of the genes in these 85 contigs did not align with the UniRef90 database, which is a lower frequency than that of known microbial genetic elements (**Figure S2B**). For these contigs, we obtained 19 clusters (Cluster 0001–0019) without singletons by clustering them with contigs that shared over 80% of their genes within a cluster and >30% average amino acid identity (**Figure 2B**). Quantification of cluster abundance through long-read mapping showed that Cluster 0001, comprising 13 contigs, exhibited the highest abundance and prevalence (**Figure 2B**). Within this cluster, 8 of the 13 contigs ranked in the top 50% in terms of all contig abundance, and one of the Cluster_0001 contig (Sample ID: OO1116B8239) was among the 20 most highly abundant contigs, suggesting that Cluster 0001 presents an abundant genetic element in human saliva (**Figure 2C**). None of the contigs of Cluster 0001 could be aligned to the plasmid or phage database with > 95% identity and > 85% coverage. These results indicate that Cluster 0001 comprises a potentially novel genetic element prevalent in the human oral microbiome.

The average contig size of Cluster 0001 was 352 kb (293–395 kb) (**Figure 2D**), and the average number of open reading frames (ORFs) was 313 (**Figure 2E**). The topology of the 13 contigs was identified as circular based on the assembly of the metaFlye^18^. Long-read mapping revealed an even sequence depth throughout the genomes, with no regions lacking mapped sequences, suggesting that these contigs are unlikely to be artificially misassembled (**Figure S3**). Notably, we observed no accumulation of alignment start and stop positions within the genome, which is a characteristic feature of linear contigs with terminal direct repeats^14^, supporting the circular topology of these contigs (**Figure S3**). Moreover, we identified the candidate replication origin and terminus via GC skew analysis (**Figure 2F**) and observed a switch in the orientation of ORFs at the replication origin and terminus (**Figure 2G** and **Figure S4**), suggesting that this Cluster is a circular replicon.

We further examined the possible reasons why these abundant genetic elements have been overlooked in previous metagenomic studies. One of the reasons could be the increased sequencing depth. Specifically, the preNuc increased the long-read sequence coverage of Cluster 0001 in a given sample from 9.6 to 30 folds, which contributed to the reconstruction of a high-quality genome (**Figure 2H**). Another reason is the challenge of assembling several regions using short reads. Although short-read assembly successfully reconstructed some parts of the contigs, these assemblies were fragmented, with 48% (48/102 points) of the breakpoints overlapping with repetitive insertion sequences (ISs) (**Figure 2I and Figure S5**). We also noted that 17% (17/102 points) of the breakpoints overlapped with other hypervariable regions (**Figure 2I and Figure S5)**. We have designated this previously unrecognized genetic element as “Inocle” (**<u>In</u>**sertion sequence encoded; **<u>o</u>**ral origin; cir**<u>cle</u>** genomic structure). Considering the lack of bacterial marker genes, Inocles are likely novel ECEs in the human oral microbiome.

### Discovery of the Inocle marker gene expands the family members of Inocle

Upon discovering the Inocle, we first investigated how many genomes within its family remain undiscovered. To this end, we searched for any marker gene that could represent an efficient strategy for identifying Inocle genomes from other metagenomic contigs. To identify such a marker gene, we clustered all Inocle ORFs and identified a single gene that satisfied the following criteria: having >70% amino acid identity and >50% alignment coverage across all initially identified Inocle genomes. We named this gene as “InoC” (Inocle conserved gene). InoC is a single-copy gene with an average amino acid length of 454 and a replication-relaxation domain (Pfam ID: PF13814) in the N-terminus domain (**Figure 3A**). No genes similar to InoC were identified in the NCBI nucleotide database, indicating that InoC is an Inocle-specific gene.

**Figure 3.**
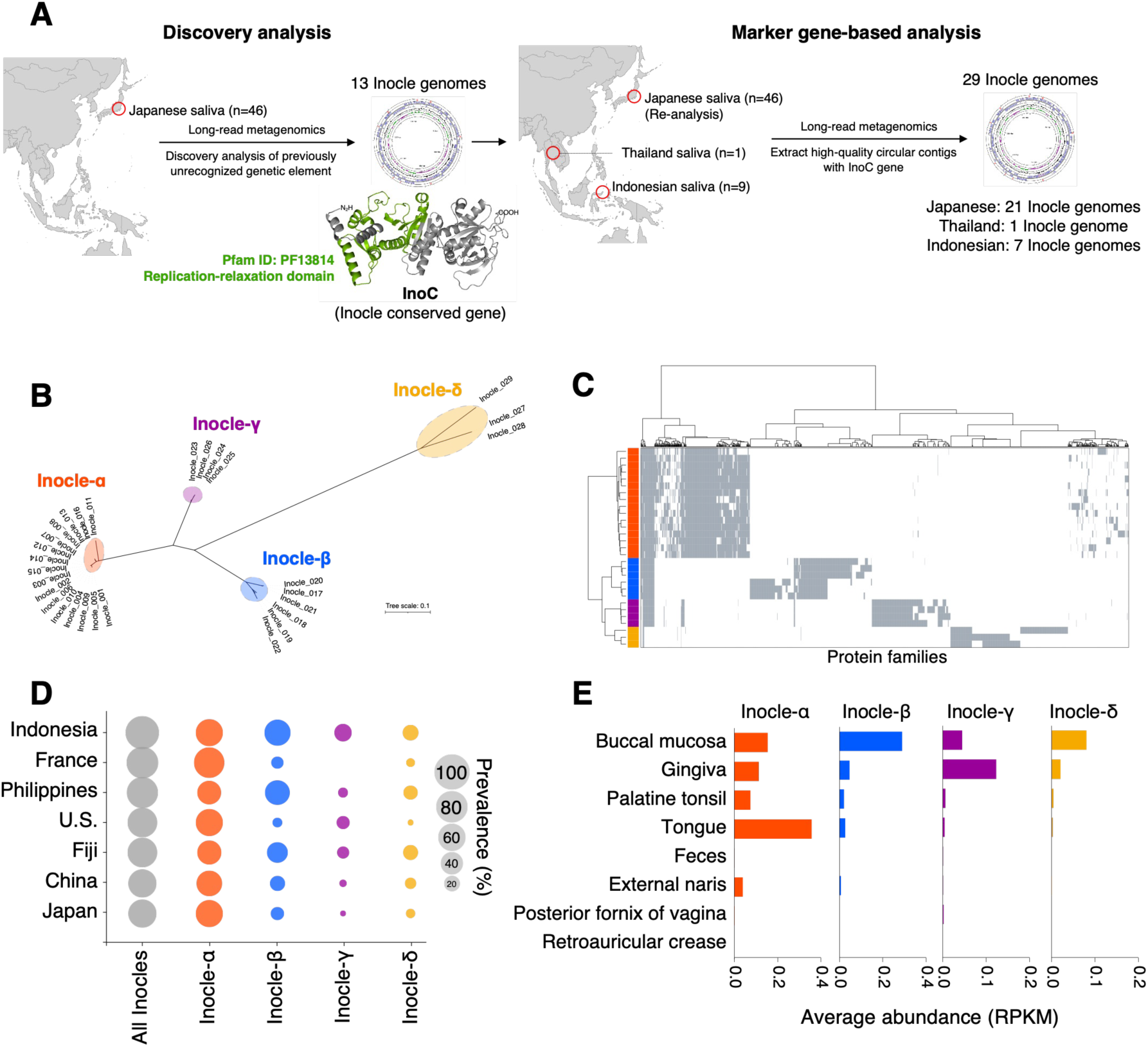
Identification and characterizations of the four Inocle taxa. (A) Workflow for identification of additional Inocle genomes. Discovery analysis using long-read contigs acquired from 46 Japanese participants identified 13 Inocle genomes and InoC genes. The representative predicted 3D structure of InoC is shown; the green region indicates the replication-relaxation domain in the InoC. To expand the number of Inocle genomes, circular contigs encoding an InoC were identified using high-quality long-read contigs obtained from 46 Japanese, 9 Indonesian, and 1 Thailand participants. (B) Phylogenetic tree of the 29 InoC amino acid sequences. (C) Comparative protein repertoires of 29 Inocle genomes. The horizontal axis indicates protein families (PFs) of Inocles, and the vertical axis represents each Inocle genome. The heatmap shows the presence (gray) or absence (white) of each PF in each genome. The taxon of each genome corresponds with the color code used in (A). (D) Prevalence of total Inocles and each Inocle taxon in the seven countries. The size of each circle shows the prevalence of each Inocle taxon in each country. (E) The average abundance of each Inocle taxon in each body site.

Using InoC, we conducted an expansion analysis to identify additional Inocle family genomes from the high-quality circular contigs, which were initially obtained from the 46 Japanese saliva samples (**Figure 3A**). To further examine the global prevalence of the Inocle family genomes, we also applied the same strategy to long-read contigs, which were assembled from saliva samples of nine Indonesian and one Thailand healthy participants (**Table S2 and S3**). We identified 16 additional circular contigs encoding InoC (**Figure 3A**), all containing candidates for replication origin and terminus (**Figure S6**). Finally, we identified 29 closed circular Inocle genomes, including 21 genomes from 17 Japanese participants, 7 genomes from four Indonesian participants, and one genome from a Thailand participant (**Table S4**).

### Discovery of four Inocle taxa and their genetical and ecological differences

The phylogenetic tree of the InoC genes of the 29 Inocle genomes revealed four distinct groups. We named these groups as Inocle-α, β, γ, and δ (**Figure 3B**). The average amino acid identity of the InoC within the same group was 94.5%, and that among different groups was 64.1% (**Figure S7A**). Inocle-α had the largest genome (368 kb on average), whereas Inocle-δ had the smallest (245 kb on average) (**Figure S7B**). The average nucleotide identity within each group was 88.9%, and we could not obtain sufficient alignment coverage (>20%) between the groups (**Figure S7C**). The number of tRNAs differed in each group (**Figure S7D**). The GC content of Inocle-α, β, and γ averaged 32%, whereas that of Inocle-δ averaged 24% (**Figure S7E**). All Inocles had IS, with Inocle-α being particularly enriched with ISs (nine ISs/genome on average), suggesting that Inocles expand their genetic diversity by the mobile genetic element of ISs (**Figure S7F)**. A total of 1,619 protein families (PFs) were identified from across the Inocle genomes, having >50% amino acid identity (**Table S5**). Among them, 1,537 PFs (95%) were only observed in a specific group (**Figure 3C**). The genetic differences between these four groups indicate that these four putative taxonomies of Inocles have different evolutionary histories.

To investigate the geographic distribution of each Inocle taxon, we downloaded salivary shotgun metagenomic sequences of healthy individuals from China, the Philippines, France, the United States, and Fiji from public databases (**Table S6**). We also obtained short-read metagenomic sequences of saliva samples from 68 healthy Japanese and 20 healthy Indonesian participants (**Table S6**). Inocle-positive samples were identified by mapping the short reads to the InoC genes, and we found that 74% of people are Inocle-positive on average (**Figure 3D**). Indonesia exhibited a higher prevalence (90%) of Inocles than other countries, whereas Japan had the lowest prevalence (64%) (**Figure 3D**). Across all countries, Inocle-α was the dominant group with 57% prevalence on average, although non-industrialized countries of Indonesia, the Philippines, and Fiji exhibited a higher prevalence of Inocle-β than that of other countries (**Figure 3D**).

Next, we mapped the shotgun metagenomic sequences of various body sites collected by the Human Microbiome Project (HMP)^19^ to the InoC genes to clarify the localization of each Inocle taxon in the human body. We detected the Inocles from oral sites, but they were rare in feces, external naris, posterior fornix of the vagina, and retroauricular crease (**Figure 3E**). Within the oral site, we revealed that the primary habitat of Inocle-α is at the tongue dorsum, that of Inocle-β and Inocle-δ is at the buccal mucosa, and that of Inocle-γ is at the gingiva (**Figure 3E**), suggesting that each taxon has a distinct habitat within the oral cavity.

### Inocles are giant plasmid-like elements of the *Streptococcus*

Bacterial cells and small extracellular particles such as phages can be separated by centrifugation followed by 0.45-μm filtration into centrifuged pellets and 0.45-μm filtered fraction, respectively^20^. We found that the number of Inocle-derived sequences was 16-fold higher in bacterial pellet fraction than in 0.45-μm filtered fraction on average after fractionating the tested saliva samples, suggesting Inocles are intracellular ECEs (**Figure 4A**). Some intracellular ECEs have homologous genes with their host bacteria^21^. Therefore, to identify the host bacterium of Inocles, we performed a homology search using the Inocle genes against all bacterial genera in the UniRef90 database. The results revealed that an average of 53 ORFs/genome were homologous to *Streptococcus* genes (**Figure 4B**), suggesting that *Streptococcus* is the candidate host bacteria of Inocles.

**Figure 4.**
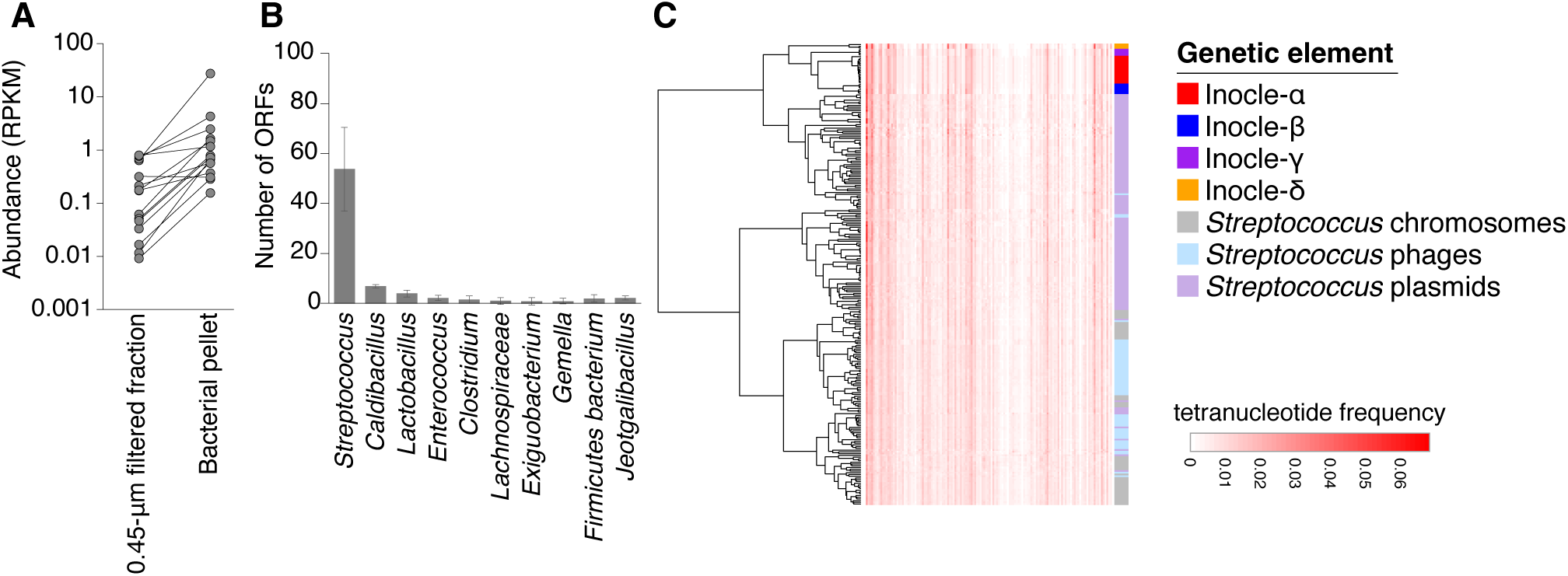
Relationships between Inocles and host bacteria. (A) Comparative abundance of Inocle genomes in the bacterial pellets and 0.45-μm filtered fraction of saliva samples. (B) The average number of Inocle ORFs aligned to genus-defined genes in the UniRef90 database. Error bars represent standard deviation (SD). (C) Genomic comparison between Inocles and known genetic elements of *Streptococcus* based on the tetranucleotide frequency.

To identify the exact host bacterial species of Inocle, we attempted to isolate *Streptococcus* strains from a saliva sample, which includes a specific Inocle-α genome (Inocle_004). Inocle-positive colonies on culture plates were detected by Sanger sequencing of PCR products specific to Inocle_004. The colony whose PCR product sequence matched perfectly with Inocle_004 was identified as the host bacterial strain of Inocle_004. The taxonomy of the host bacterium was determined as *Streptococcus salivarius* based on whole genome shotgun sequencing. We obtained sequences with more than 2,500 sequence depth of this isolate but which did not include Inocle-derived sequences, suggesting the ratio of host bacterium and Inocle_004 is <1/2500 under culturing conditions. On the other hand, the ratio of Inocle to host bacteria in native saliva was 1:15.

A comparison of genomic similarity based on tetranucleotide frequency showed that Inocles have a distinct nucleotide composition that is clearly different from the known genetic elements of *Streptococcus* but similar to some plasmids compared with chromosomes and phages (**Figure 4C**). We identified the VirB4 and VirD4 in all Inocle genomes (**Table S5**), which are required to transfer plasmid from donor to recipient cells via the type IV secretion system^22^. Hence, Inocles are considered novel giant plasmid-like elements of *Streptococcus*, possibly having the ability for horizontal transmission.

### Gene annotation of the Inocle genomes

To reveal the biological roles of Inocles in host bacteria, we conducted functional annotation of 1,619 PFs encoded by the Inocle genomes (**Table S5**). First, we considered the subcellular localization of these PFs and identified 644 (40%) and 620 (38%) PFs as cytoplasmic and unknown subcellularly localized proteins, respectively (**Figure 5A**). Among these 644 cytoplasmic proteins, we annotated 21 functional genes using Prokka^23^. Among the 11 conserved cytoplasmic genes in all Inocle taxa, 6 were annotated as genes associated with DNA repair and recombination (ko03400) based on the Kyoto Encyclopedia of Genes and Genome (KEGG) BRITE database. All these genes contained DNA damage repair-related functions, such as those of exonuclease, DNA gyrase, and DNA polymerase III (**Figure 5B**). We identified a RecD-like DNA helicase with unknown subcellular localization genes, which was also related to DNA damage repair-related genes (**Figure 5C**). We found other stress response-related genes detected in multiple taxa, such as RpoS contributing stress responses against multiple stressors^24^ and NrdH conferring oxidative stress tolerance^25^. We confirmed the transcriptional activity for six of seven DNA damage repair genes, RpoS, and NrdH (**Figure 5B and C**), based on the metatranscriptome sequences of human saliva^26^.

**Figure 5.**
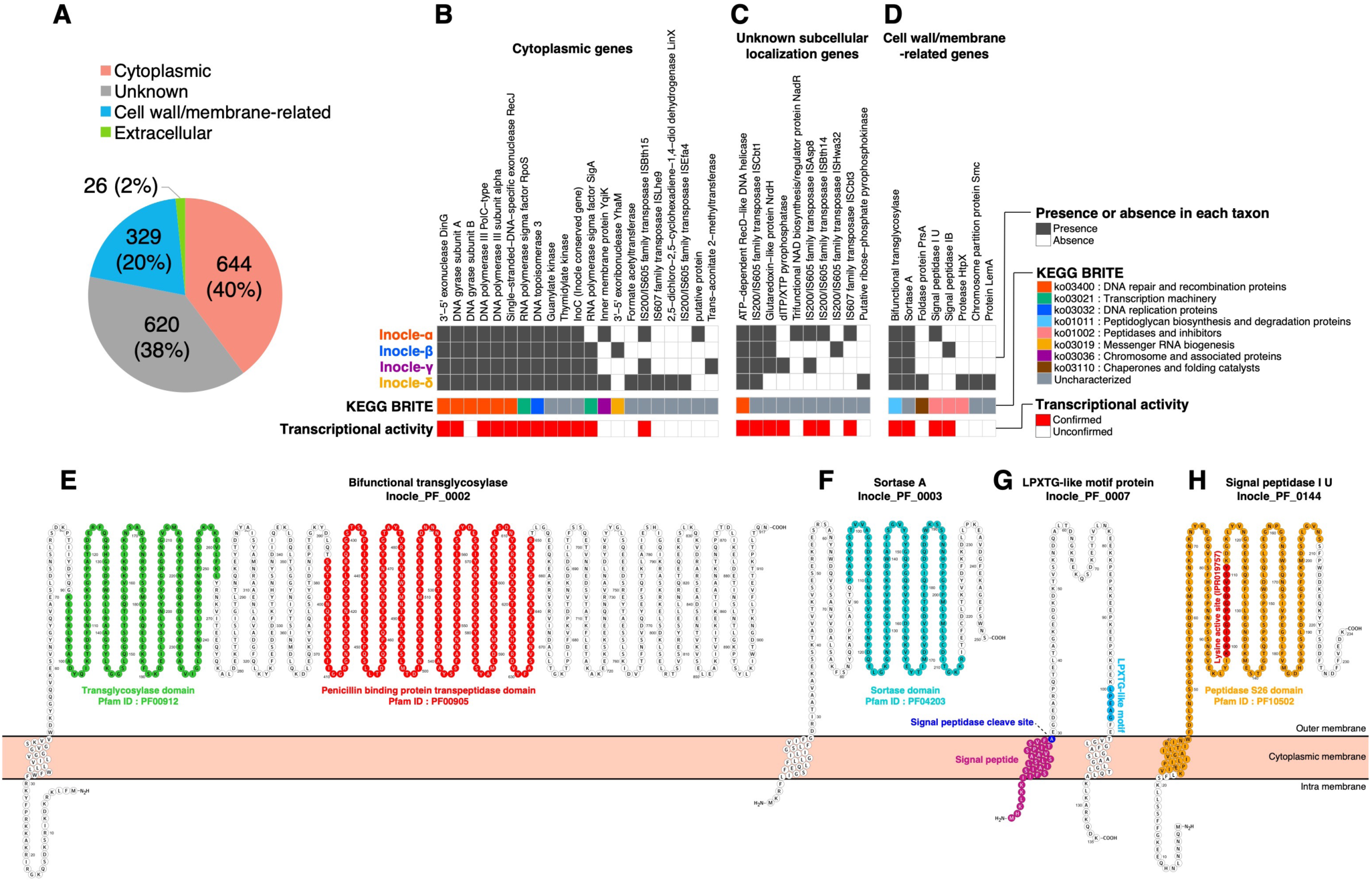
Functional characterization of Inocles. (A) The ratio of subcellular localization in 1,619 Inocle protein families. (B–D) Functionally annotated genes of Inocles in cytoplasmic genes (B), unknown subcellular localization genes (C), and cell wall/membrane-related genes (D). The gray and white heatmap indicates the existence of each gene in each Inocle taxon. The functional categories of the KEGG BRITE are shown for each gene. Genes with confirmed transcriptional activity in saliva are shown in red heatmap. (E–H) Representative 2D structure of the bifunctional transglycosylase (E), sortase A (F), LPXTG-like motif protein (G), and signal peptidase (H) on the membrane.

Next, we focused on 329 PFs (20%) of possible cell wall/membrane-related proteins (**Figure 5A**). Among them, we identified eight functionally annotated genes, two of which were conserved across all Inocle taxa, while one was classified as peptidoglycan biosynthesis/degradation protein based on the KEGG BRITE database (**Figure 5D**). This protein comprises a transmembrane domain at its N-terminus as well as a transglycosylase domain and a penicillin-binding protein transpeptidase domain in the outer membrane region (**Figure 5E**). Therefore, it is a potential bifunctional transglycosylase that catalyzes the polymerization and cross-linking of the glycan chain for peptidoglycan biosynthesis^27^. The other conserved cell wall/membrane-related genes were annotated as sortase A (**Figure 5D**), which comprises a transmembrane domain at its N-terminus, with the sortase domain located in the outer membrane region (**Figure 5F**). This protein possibly recognizes the LPXTG motif in cell wall proteins and anchors them to peptidoglycan^28^. We identified six LPXTG-like motif PFs presumed to be recognized by sortase A. All Inocle taxa encoded at least one LPXTG-like motif PF (**Figure S8A**). These LPXTG-like motif PFs contain two transmembrane domains, one at the N-terminus and one at the C-terminus, spanning the outer membrane region (**Figure 5G and Figure S8B)**. We found that an LPXTG-like motif is in the outer membrane region of the C-terminus, with the signal peptide for membrane transition and the signal peptidase cleavage position located at the N-terminus (**Figure 5G and Figure S8B**). We identified signal peptidases from Inocle-α, β, and γ with the Signal peptidase S26 domain (PF10502) (**Figure S8C**) and a lysine-active site for cleaving signal peptides (**Figure 5H and Figure S8D**). We confirmed the transcriptional activity for these cell wall-related genes, suggesting their bioactivity in a native oral environment (**Figure 5D, Figure S8A, and S8C**).

### Inocle-α is correlated with human systemic immune statuses

We then investigated the associations of Inocles with human physiologies. We first examined whether Inocle-α is associated with the sex and age of the host human (healthy participants) and observed no significant associations (**Table S7, Figure S9A, and S9B**). Owing to the low prevalence of Inocle-β, γ, and δ in our Japanese sample population, corresponding data could not be statistically analyzed (**Figure 3E**).

We further examined the associations using a multi-omics dataset of humans. We collected peripheral blood mononuclear cells (PBMCs) from the 29 individuals whose saliva was collected in this study and conducted the single-cell RNA sequencing (scRNA-seq) analysis (**Table S8**). The cellular populations of PBMCs were annotated into 19 cell types based on Azimuth references^29^. Correlation analysis revealed a significantly positive correlation of Inocle-α with intermediate and memory B cells but not with plasma cells (**Figure 6A**). We also observed a significantly negative correlation between Inocle-α and CD16+ monocytes (**Figure 6A**). These associations were not observed for *S. salivarius* (**Figure 6A**). Comparison between the Inocle-α-positive (n = 19) and - negative (n = 10) groups revealed that intermediate and memory B cells were significantly increased in the Inocle-α-positive group, whereas CD16+ monocytes were significantly decreased compared to those in the Inocle-α-negative group (**Figure 6B**).

**Figure 6.**
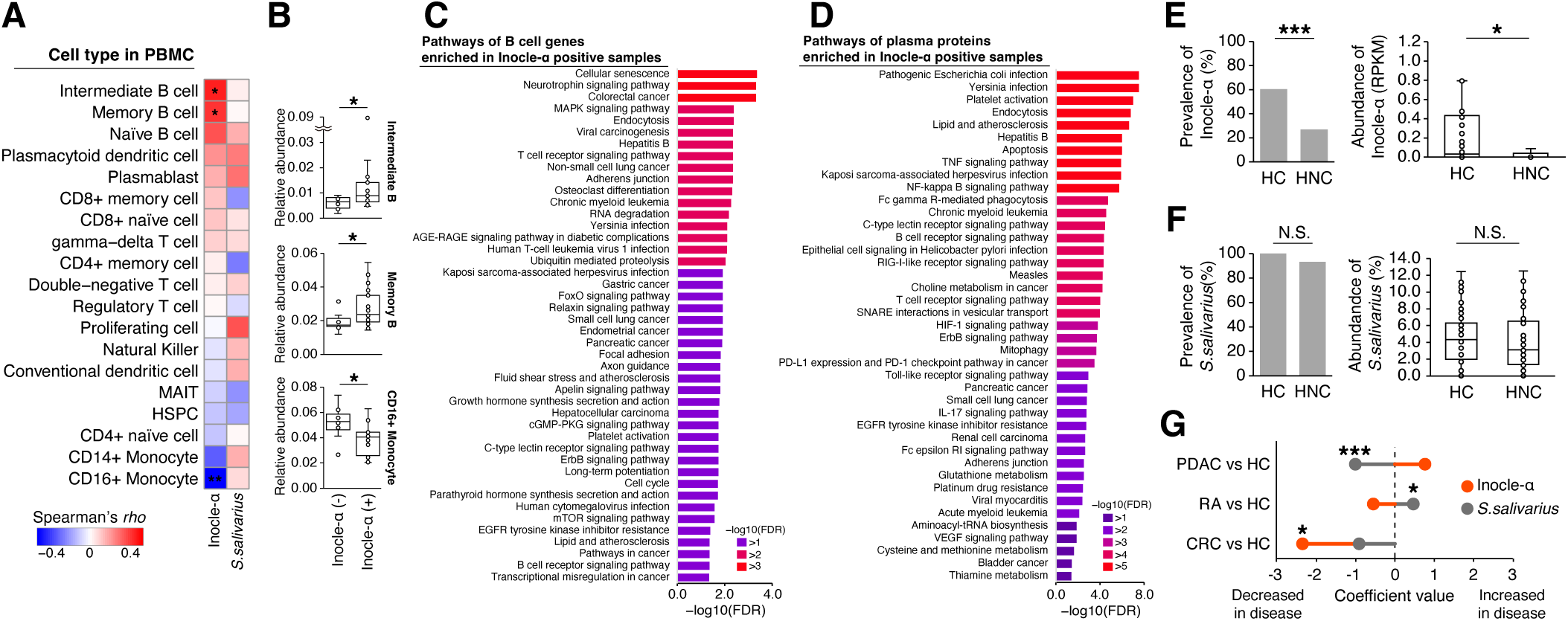
Association of Inocle-α with human physiologies. (A) Spearman’s *rho* between PBMC populations and the abundance of Inocle-α and *S. salivarius.* \**p* < 0.05, \*\**p* < 0.01. (B) Comparison of the relative abundance of intermediate B cells, memory B cells, and CD16+ monocytes between the Inocle-α-positive and -negative groups. \**p* < 0.05. Statistical results were obtained using the Wilcoxon rank-sum test. (C) Gene Ontology (GO) enrichment analysis of significantly increased gene expression of B cells in the Inocle-α-positive group compared to the Inocle-α-negative group. (D) GO enrichment analysis of significantly increased plasma proteins in the Inocle-α-positive group compared to the Inocle-α-negative group. (E) and (F) Comparison of the prevalence and abundance of Inocle-α (E) and *S. salivarius* (F) between healthy controls (HCs) and patients with head and neck cancer (HNC). The bar plots indicate the prevalence of Inocle-α and *S. salivairus* in the HC and HNC groups. The box plots indicate the abundance of Inocle-α and *S. salivarius* in the HC and HNC groups. The outliners are removed for visualization in box plots. \**p* < 0.05, \*\*\**p* < 0.001, N.S., not significant. Statistical results were obtained using the Fisher’s exact test for prevalence and the MaAsLin2 for abundance. (G) Comparison of the abundance of Inocle-α and *S. salivarius* between patients with different diseases and HCs in each study. The coefficient values and *p*-value calculated via MaAsLin2 are shown. \**p* < 0.05. Abbreviations; HSPC: Hematopoietic stem and progenitor cell, MAIT: Mucosal associated invariant T, CRC: colorectal cancer, PDAC: pancreatic ductal adenocarcinoma, RA: rheumatoid arthritis.

Given the positive correlations between several B cells and Inocle-α, we further analyzed gene expression in B cells. We observed that the expression of 687 and 62 genes was significantly increased and decreased in the Inocle-α-positive group compared to the Inocle-α-negative group, respectively (**Table S9**). Gene ontology (GO) analysis revealed 44 enriched pathways in the Inocle-α-positive group compared to the Inocle-α-negative group (>1.4 -log10 FDR [<0.05 FDR]) (**Figure 6C**). Among these 44 pathways, cellular senescence was the most enriched pathway in the Inocle-α-positive group (**Figure 6C**). Moreover, five pathways were involved in the response to bacterial and viral infections, including hepatitis B, Yersinia infection, human T-cell leukemia virus 1 infection, Kaposi sarcoma-associated herpesvirus infection, and human cytomegalovirus infection (**Figure 6C**).

To further scrutinize the associations between Inocle-α and human systemic immune systems, we analyzed the expression data of a total of 5,420 plasma proteins using proximity extension assay technology (Olink) from 40 healthy Japanese participants, including 27 Inocle-α-positive and 13 Inocle-α-negative participants (**Table S10**). Comparison analysis detected 811 and 10 significantly increased and decreased proteins in the Inocle-α-positive group compared to the Inocle-α-negative group, respectively (**Table S11**). GO enrichment analysis revealed that ten pathways were particularly enriched in the Inocle-α-positive group (>5 -log10 FDR), four of which were related to the response to bacterial and viral infections, including pathogenic Escherichia coli infection, Yersinia infection, hepatitis B, and Kaposi sarcoma-associated herpesvirus infection (**Figure 6D**), partially consistent with the transcriptional differences observed in B cell populations (**Figure 6C**). In addition, we noted higher expression levels of proteins related to induction of Platelet activation, Endocytosis, Lipid and atherosclerosis, Apoptosis, TNF signaling, and NF-κB signaling pathway in the Inocle-α-positive group (**Figure 6D**).

### Inocle-α is negatively associated with patients of several cancer types

Notably, we found that Inocle-α was associated with several cancer-related pathways in the plasma proteome. These pathways included Choline metabolism in cancer, PD-L1 expression and PD-1 checkpoint pathway in cancer, pancreatic cancer, small cell lung cancer, and bladder cancer (**Figure 6D**). Therefore, we subsequently investigated the possible association between Inocle-α and cancers. We collected saliva samples from 45 patients with head and neck cancer (HNC), including 22 treatment-naïve patients, and obtained metagenomic short reads (**Table S7**). We compared the abundance of Inocle-α among 53 healthy controls (HCs) and 45 patients with HNC while considering their ages, sexes, smoking status, and cancer treatment information as possible confounding factors (**Table S7**). The prevalence of Inocle-α was significantly decreased in the HNC group (27%) compared to that in the HC group (60%), and correspondingly, the abundance of Inocle-α was decreased (**Figure 6E**). However, we could not observe a significant reduction in the prevalence and abundance of *S. salivarius* in the HNC group compared to that in the HC group (**Figure 6F**). These results suggest that host bacteria harboring Inocle-α are selectively decreased in the saliva of patients with HNC.

We further analyzed whether the associations of Inocles could be observed in other cancer types. To this end, we downloaded public salivary metagenomic short reads of patients with pancreatic ductal adenocarcinoma (PDAC)^30^ and those with colorectal cancer (CRC)^31^, as well as HCs from each study (**Table S12**). As a reference for an inflammatory disease, the data of patients with rheumatoid arthritis (RA)^32^ were also obtained (**Table S12**). Statistical analysis revealed that the abundance of *S. salivarius* was significantly altered in PDAC and RA, but that of Inocle-α was not altered (**Figure 6G**). In contrast, Inocle-α was significantly decreased in the CRC group compared to the HC group (**Figure 6G**), suggesting that the features of Inocle-α are specifically associated with disorders occurring in the gastrointestinal tract.

## DISCUSSION

Despite Inocles spreading worldwide, they have not been previously identified nor included in reference genomes, partially due to their harboring ISs or hypervariable regions. The low score of the plasmid prediction tool for Inocles reflects the difficulty in detecting Inocles using current bioinformatics tools. These results demonstrate the importance of long-read metagenomics technology in a reference-independent strategy for identifying ECEs.

The biggest feature of Inocles is their large genome size, with a maximum genome size of 395kb, possibly being one of the largest ECEs in the human commensal bacteria. We speculated that Inocles are plasmid-like elements based on nucleotide composition, suggesting that Inocles might be able to be categorized as megaplasmids^33^. Megaplasmids have been found in several human pathogens, and they carry multiple antimicrobial resistance genes to confer multi-drug resistance and increase the infectious ability^34,35^. On the other hand, Inocles does not contain antibacterial or virulent genes, suggesting that Inocles provide different benefits to the host bacteria than to the previously identified human-associated megaplasmids.

We identified ORFs of Inocles encode a variety of stress response proteins. For example, RpoS is a sigma factor that globally regulates bacterial transcription and induces stress responses, facilitating survival during environmental stresses^24^. Notably, we identified a series of genes related to the DNA damage repair pathway from Inocles, including DNA polymerase III, gyrase, RecD, RecJ, and DinG, which are activated by oxidative stress^36–39^. Additionally, we identified the peroxidase cofactor (NrdH) in Inocle-α, β, and γ, which reduces intracellular reactive oxygen species (ROS) levels^25^. These results suggest that Inocles is a key factor in resisting harsh environments, including under oxidative stresses and DNA damage in the oral environment (**Figure S10**).

Another functional feature of Inocle is that 20% of all genes in Inocles are involved in cell wall/membrane-related functions. The discovery of a bifunctional transglycosylase^27^, signal peptidase, sortase A, and LPXTG-like motif protein suggests the following processes on the cytoplasmic membrane. First, the bifunctional transglycosylase of Inocles promotes peptidoglycan biosynthesis in host bacteria^27^. Following translation, LPXTG-like motif proteins are transported to the cytoplasmic membrane, where the signal peptidase cleaves the N-terminal signal peptide of LPXTG-like motif proteins^40^. Sortase A then cleaves the LPXTG-like motif and anchors matured protein to peptidoglycans as a cell surface protein^41^. Several studies on biofilms on human epithelial cells have revealed that sortase A contributes to bacterial adhesion and evasion of the human immune system^28,42,43^. Although the functional roles of LPXTG-like motif proteins are uncharacterized, our findings suggest that cell wall-associated proteins of Inocles are involved in the colonization of *Streptococcus* to human epithelial cells.

One of the highlights of this study is the elucidation of the association between Inocle-α and human physiology. We observed a clear positive correlation between the Inocle-α and signaling responses against infectious pathogens based on the scRNA-seq and plasma proteome dataset (**Figure S10**). Although this study does not provide mechanistic insights into the interactions between Inocles, host bacteria, and the human immune system, these results suggest that Inocle-α is involved in latent or tolerant immune responses. Another notable finding from our study is the disease-specific reduction in Inocle-α. Specifically, we observed a reduction in Inocle-α in HNC and CRC but not in PDAC and RA. One plausible explanation of these observations is that alterations in the immune conditions varying across cancer types^44^ change the necessity of Inocles for *Streptococcus* to adapt to immune responses. Another possibility is that Inocle-positive individuals are less likely to develop the corresponding cancers owing to their predisposed immune conditions being unfavorable for cancer development. In either case, Inocle has the potential to serve as a biomarker for a non-invasive test for those cancers (**Figure S10**).

The biological roles of Inocles remain largely unknown, as the functions of 95% of their genes remain elusive, but the identification of *S. salivarius*, the specific host bacterium of Inocles, is probably able to verify its functions experimentally. Overall, despite only providing introductory findings, this study is anticipated to pave the way for further understanding the multi-faceted role or biological relevance of Inocles in modulating the adaptive capacity of oral bacteria and potentially impacting human health.

## ACKNOWLEDGMENTS

The authors thank M. Tsubaki, S. Imamura, and K. Abe for technical support in the experiments. The authors thank E. Ishikawa, R. Fujinaga, and S. Takashima for technical support in informatics. The authors thank Prof. Dr. dr. Nova H. Kapantow, DAN, M.Sc, Sp.GK for the saliva sample collection from Indonesian participants. The authors thank K. Matsuura for collecting saliva samples from patients with HNC. The authors thank S. Stamouli and D. Takewaki for the critical reading of this manuscript. The authors thank M. Iwanaga for technical support in the isolation of *Streptococcus*. This study was supported by grants from the Institute for Fermentation, Osaka (Y-2022-1-010), the Japan Agency for Medical Research and Development (AMED) (22fk0108538s0201), and the JSPS KAKENHI (24K18092) to Y.Kiguchi, a grant from the JSPS KAKENHI (22K15833) to T.E., and a grant from the Platform for Advanced Genome Science (22H04925) to Y.S.

## AUTHOR CONTRIBUTIONS

Y.Kiguchi and Y.S. conceived and supervised this study. Y.S. managed sequencing and bioinformatics platforms. N.H., Y.Kiguchi, and T.M. developed the preNuc method. Y.Kiguchi, N.H., and T.M. contributed to the metagenomic DNA extraction, sequencing, and analysis. L.R.R., J.S.B.T., Y.Kiguchi, and T.M. collected saliva samples from Indonesian participants. A.I. and T.M. isolated the *Streptococcus* strain with Inocle. T.E., S.K., T.F., N.T., M.T., and R.Y. collected the saliva samples from patients with HNC. Y.Kashima and A.K. obtained sequencing data for scRNA-seq and the Olink proteome from the blood samples. Y.Kashima analyzed the scRNA-seq dataset. Y.Kiguchi analyzed the Olink proteome dataset. Y.Kuze developed the bioinformatics environment. Y.Kiguchi, T.M., and Y.S. wrote the manuscript. All authors have read and approved the manuscript.

## SUPPLEMENTAL INFORMATION

**Figure S1.**
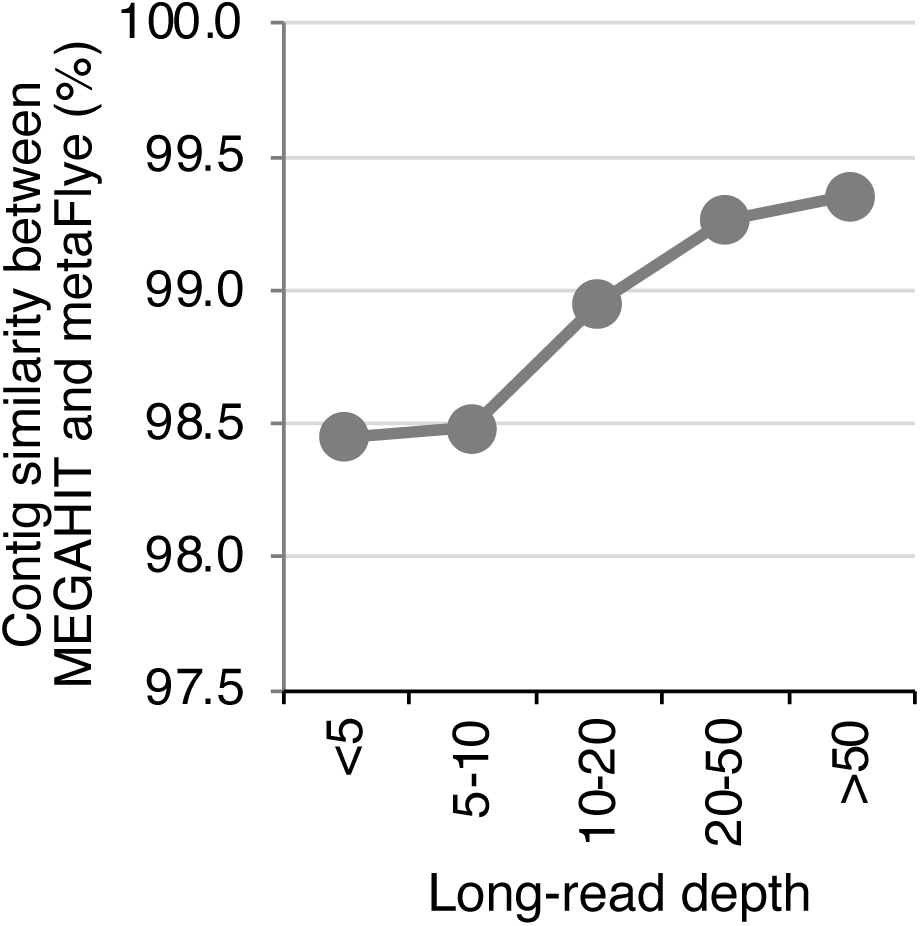
Sequence similarity between long-read contigs and short-read contigs. Sequence similarity between PromethION and NovaSeq contigs acquired from the same samples (sample ID; m_5 and k_5) assembled using metaFlye and MEGAHIT. The y-axis indicates the average sequence similarity of the PromethION contigs with the best-hit NovaSeq contigs corresponding to each long-read depth of the PromethION contigs (x-axis).

**Figure S2.**
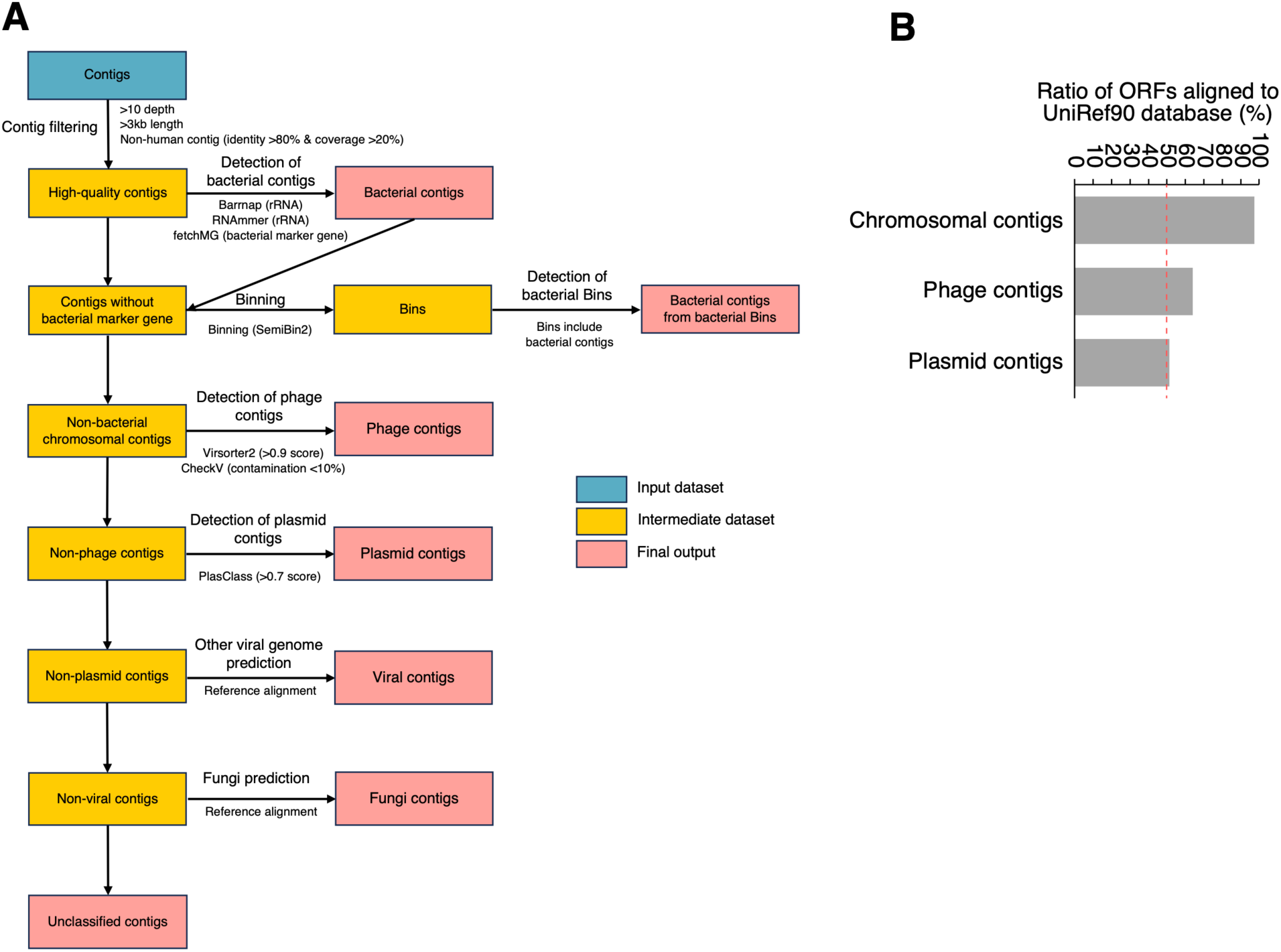
Workflow for the classification of long-read contigs as microbial genetic elements. (A) The flowchart shows the pipeline for classifying long-read contigs as each genetic element. Human genome-derived, low-quality (<10 depth), and small contigs (<3kb) were removed. Subsequently, bacterial chromosomal contigs were identified based on the presence of bacterial marker genes of rRNA and fetchMG-defined marker genes. Contigs belonging to the same Bin, including bacterial chromosomal contigs, were identified as potential bacterial chromosomal contigs. Non-bacterial chromosomal contigs satisfying >0.9 virsorter2 score and <10% contamination (CheckV) were identified as phage contigs. Among non-phage contigs, plasmid contigs (score >0.7) were identified using PlasClass. Eukaryotic viral and fungal contigs were identified based on their alignment with reference genomes. Finally, contigs that were not identified as genetic elements were considered to be unclassified contigs. (B) The ratio of open reading frames (ORFs) aligned to the UniRef90 database in all ORFs of each known genetic element.

**Figure S3.**
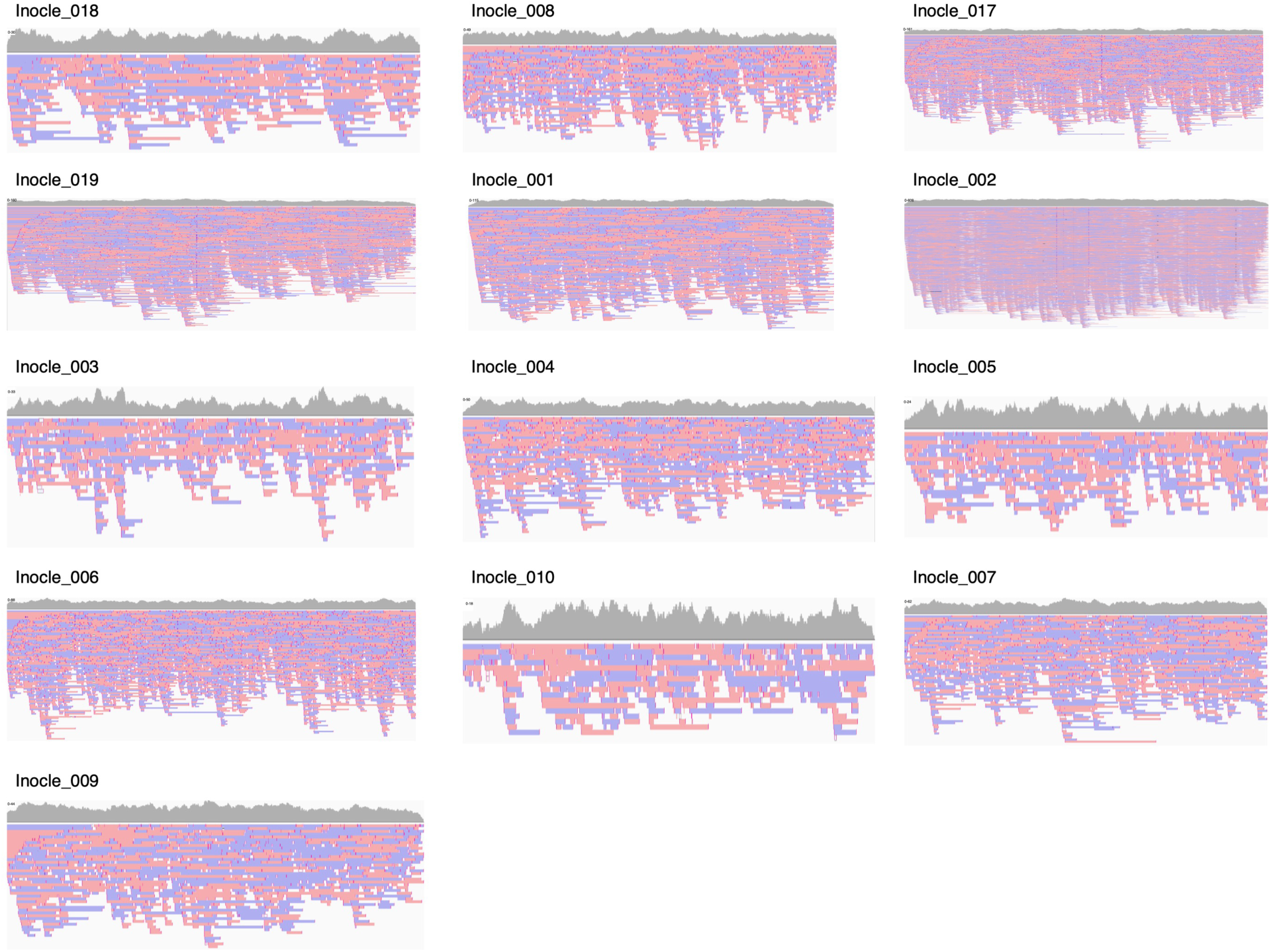
Long-read mapping results of the 13 contigs belonging to Cluster 0001. The gray bar plot in the mapping results of the Integrative Genomics Viewer (IGV) indicates the depth of the mapped long reads. Forward and reverse reads are displayed in pastel red and blue boxes, respectively.

**Figure S4.**
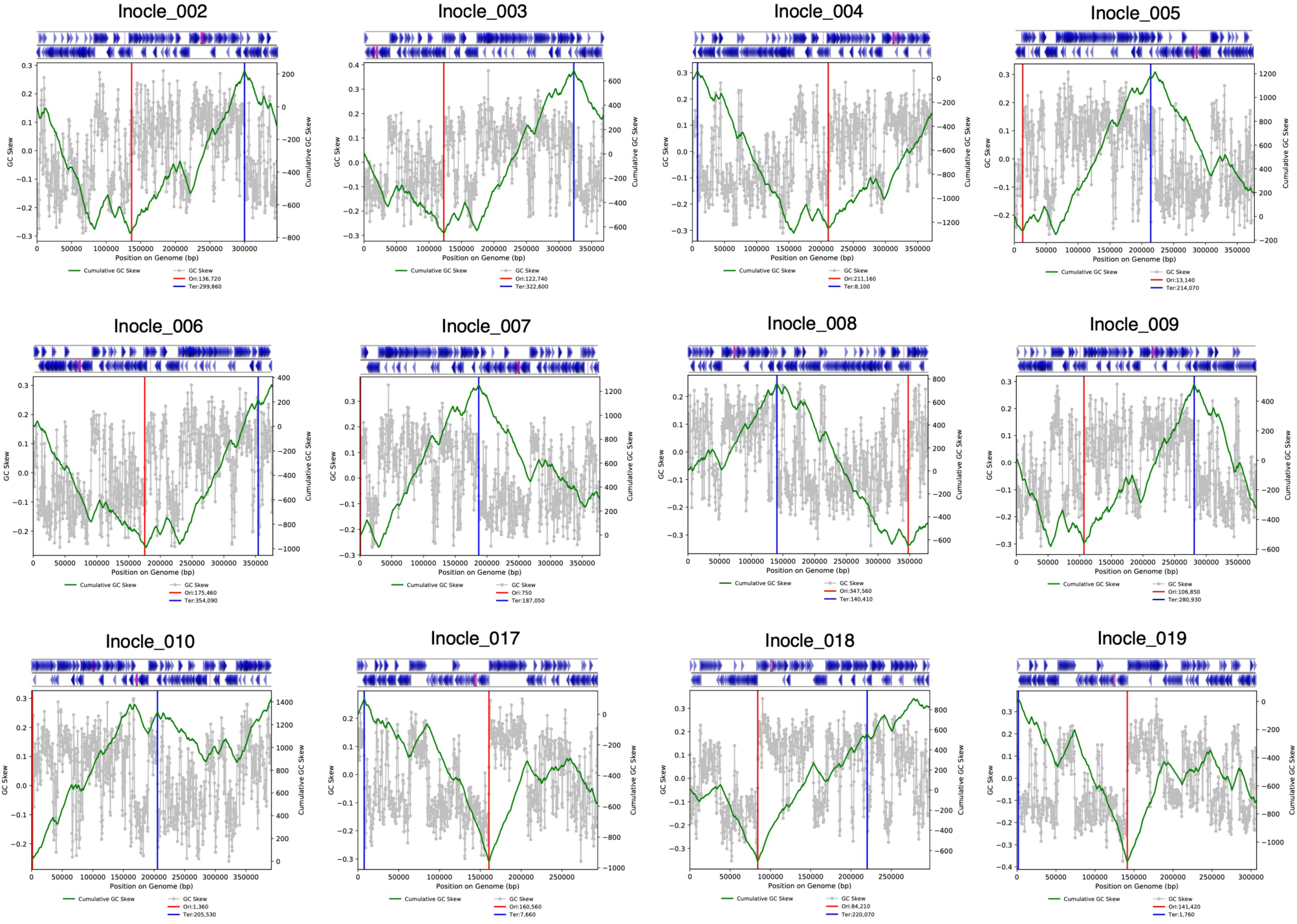
Replichore structures in 13 contigs belonging to Cluster 0001. The green and gray lines indicate the cumulative GC skew and the GC skew at each genomic position, respectively. The candidate replication origins and termini are indicated using vertical red and blue lines, respectively. The blue arrows in the upper figures represent the forward strand and the reverse strand of ORFs.

**Figure S5.**
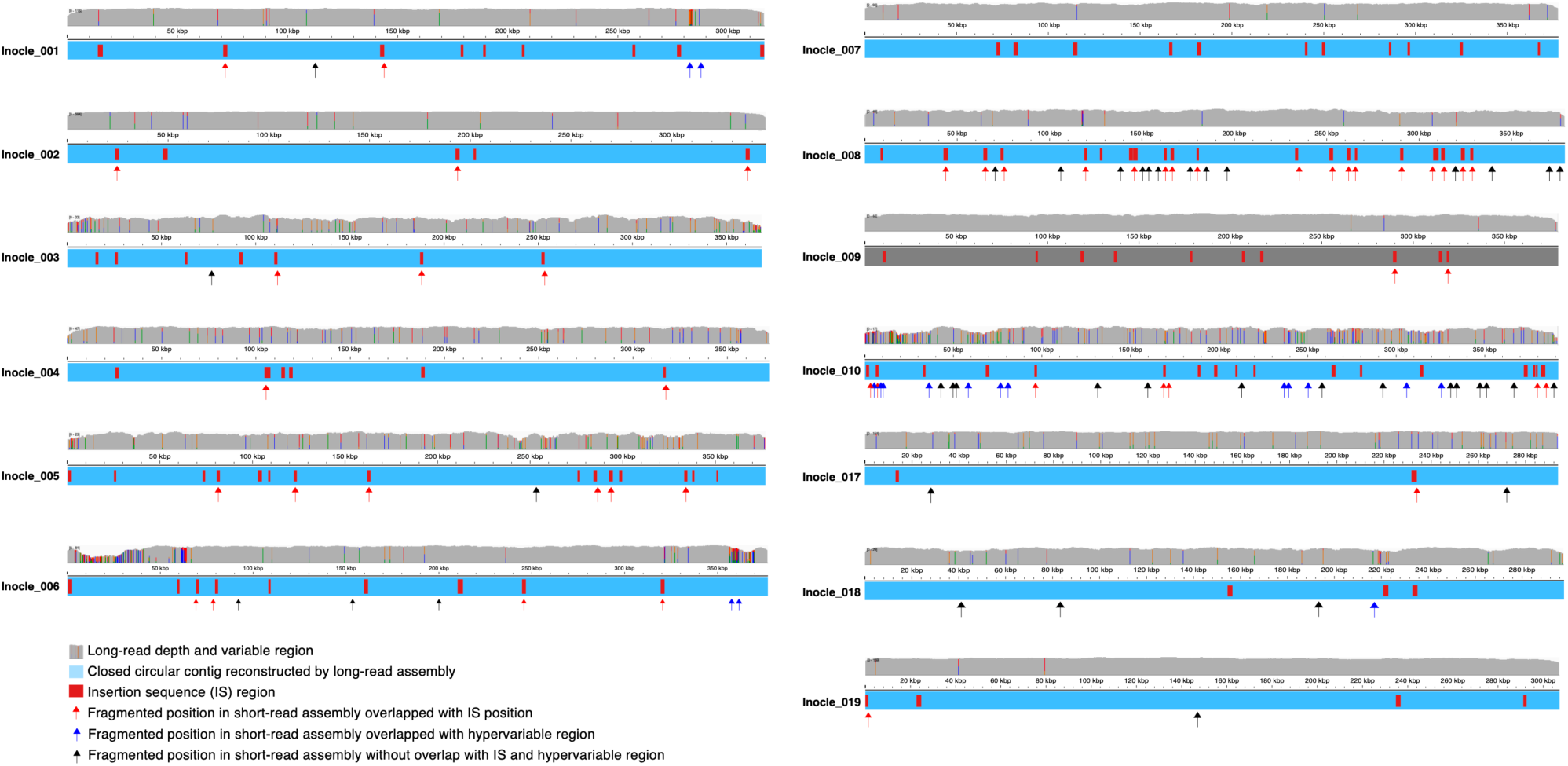
Impact of the long-read metagenomics for reconstructing of Inocle genomes. Comparison of the long-read and corresponding short-read contigs. The blue bar represents the complete genome constructed through long-read assembly. The red boxes indicate the insertion sequence (IS) regions in the contig. Arrows represent the fragmented positions in the short-read assembly. The upper gray bar plot indicates the depth of the mapped long reads. Colored bars in the depth bar plots indicate single-nucleotide variants: Green=A, red=T, orange=G, and blue=C.

**Figure S6.**
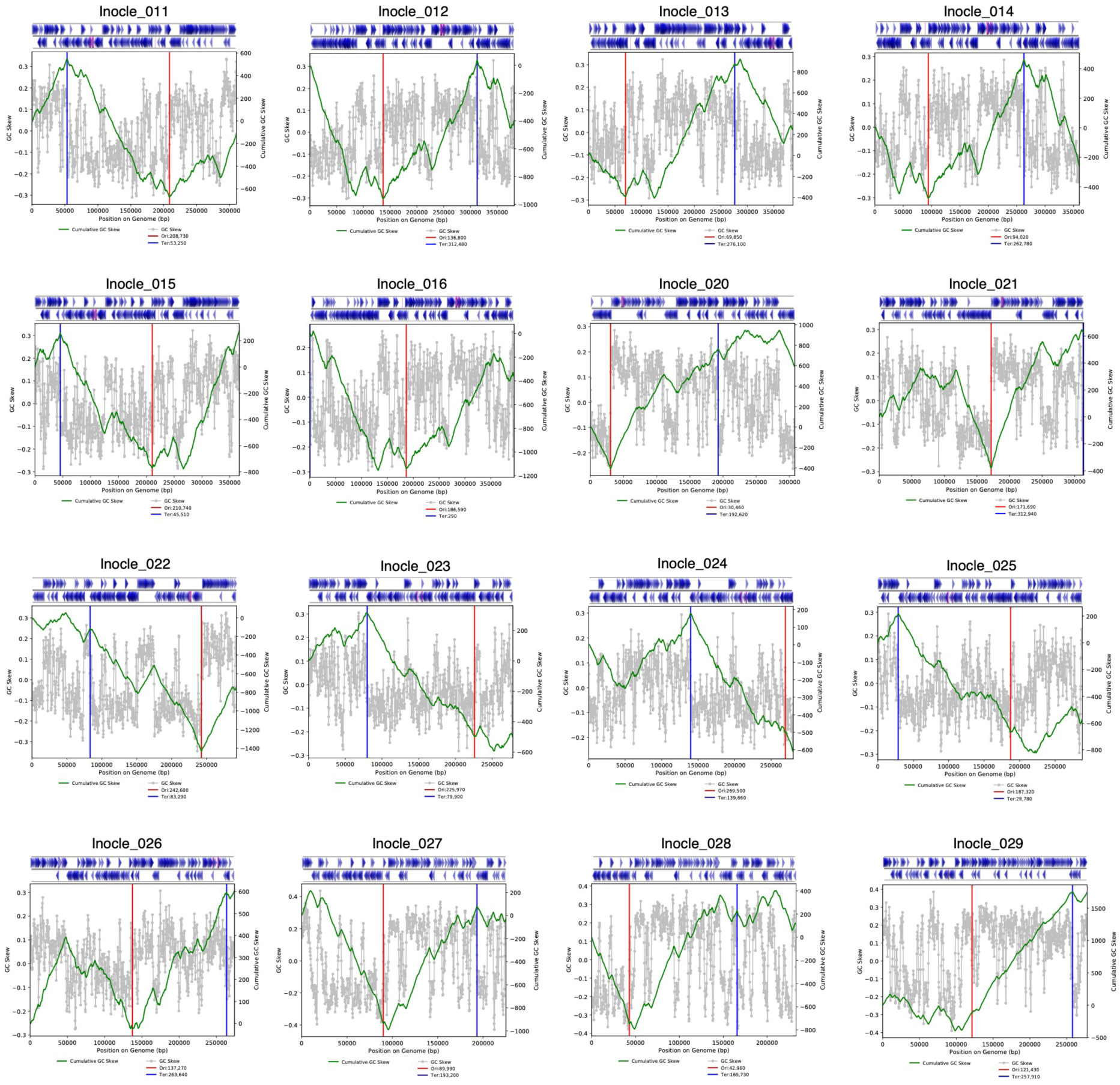
Replichore structures in 16 additional Inocle genomes. The green line indicates the cumulative GC skew and the gray plots indicate the GC skew at each genomic position. The candidate replication origins and termini are indicated by vertical red and blue lines, respectively. The blue arrows in the upper figures represent the forward strand and the reverse strand of ORFs.

**Figure S7.**
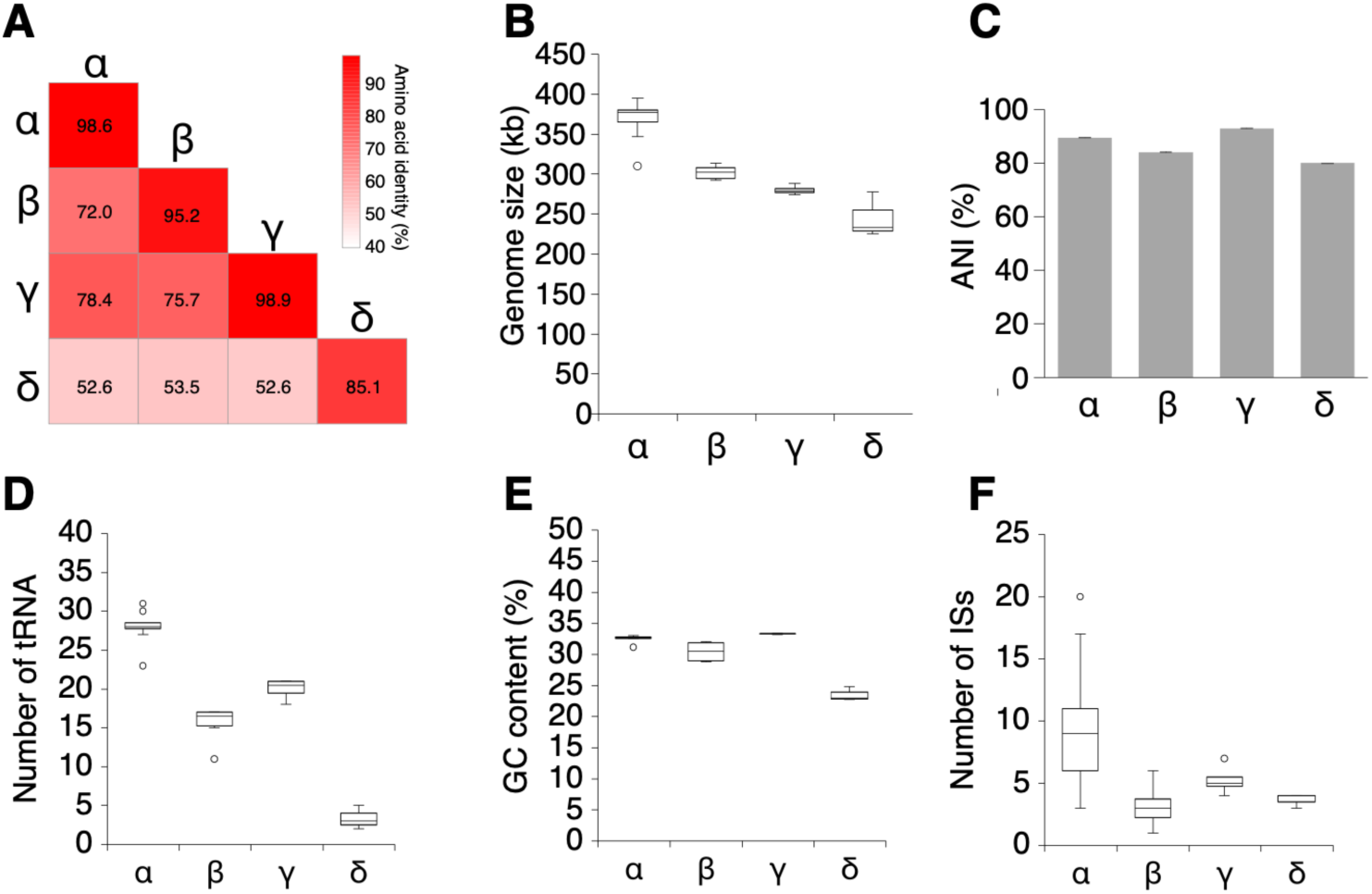
Genomic differences between the four Inocle taxa. (A) Amino acid identity of the InoC gene among Inocle taxa. The heatmap indicates the average amino acid identity (AAI) of the InoC gene in each pair of comparisons, and the numbers in the heatmap indicate AAI. (B) The distribution of the genome size and (C) average nucleotide identity in each Inocle taxa; error bars represent standard deviation (SD). (D) The number of tRNAs, (E) GC content, and (F) the number of insertion sequences (ISs) in each Inocle taxa. (B–F) Box plots represent the inter-quartile range (IQR), and lines inside the box indicate the median. Whiskers show 1.5 IQR.

**Figure S8.**
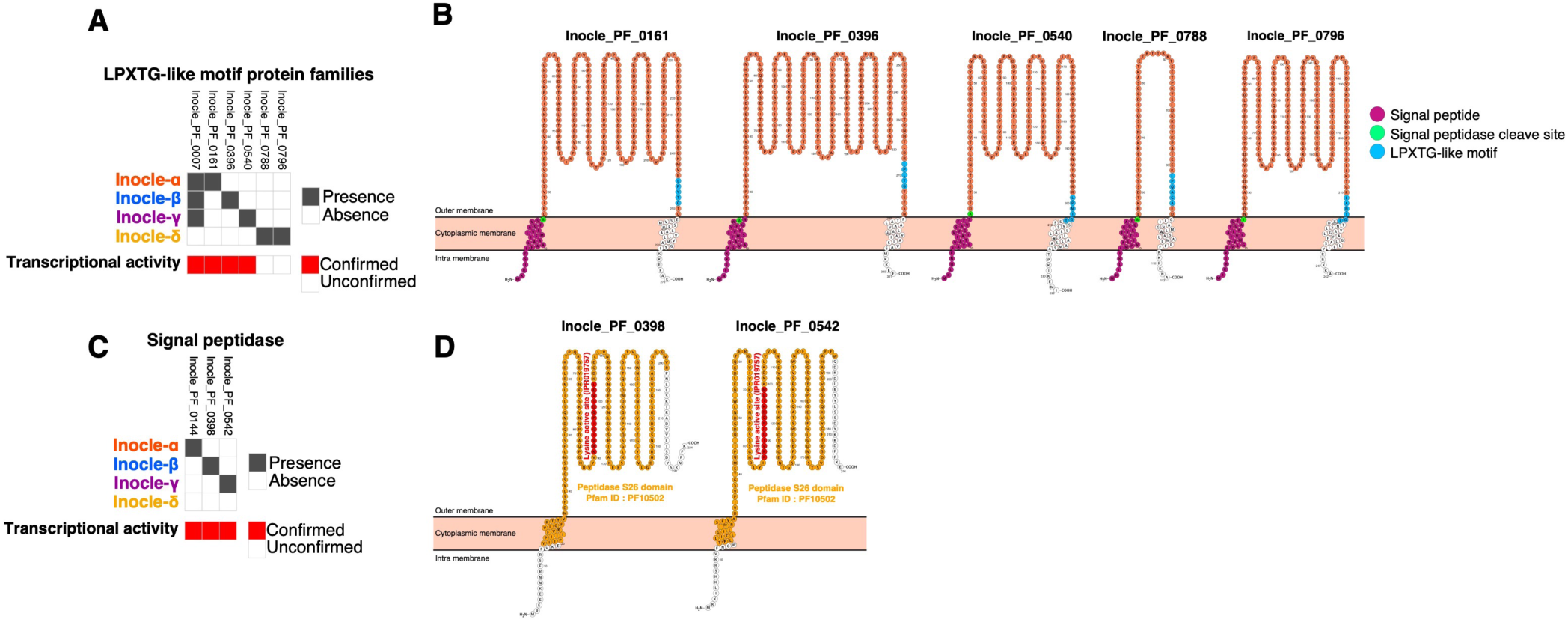
Functional annotation of the cell wall-related proteins of Inocles. (A) Presence or absence of PFs with LPXTG-like motif in each Inocle taxa. Genes with confirmed transcriptional activity in saliva are shown in red heatmaps. (B) The 2D structure of the LPXTG-like protein motif on the membrane. (C) Presence or absence of PFs with Signal peptidase S26 domain (PF10502) in each Inocle taxa. Genes with confirmed transcriptional activity in saliva are shown in red heatmaps. (D) The 2D structure of the signal peptidase (Inocle_PF_0398) on the membrane.

**Figure S9.**
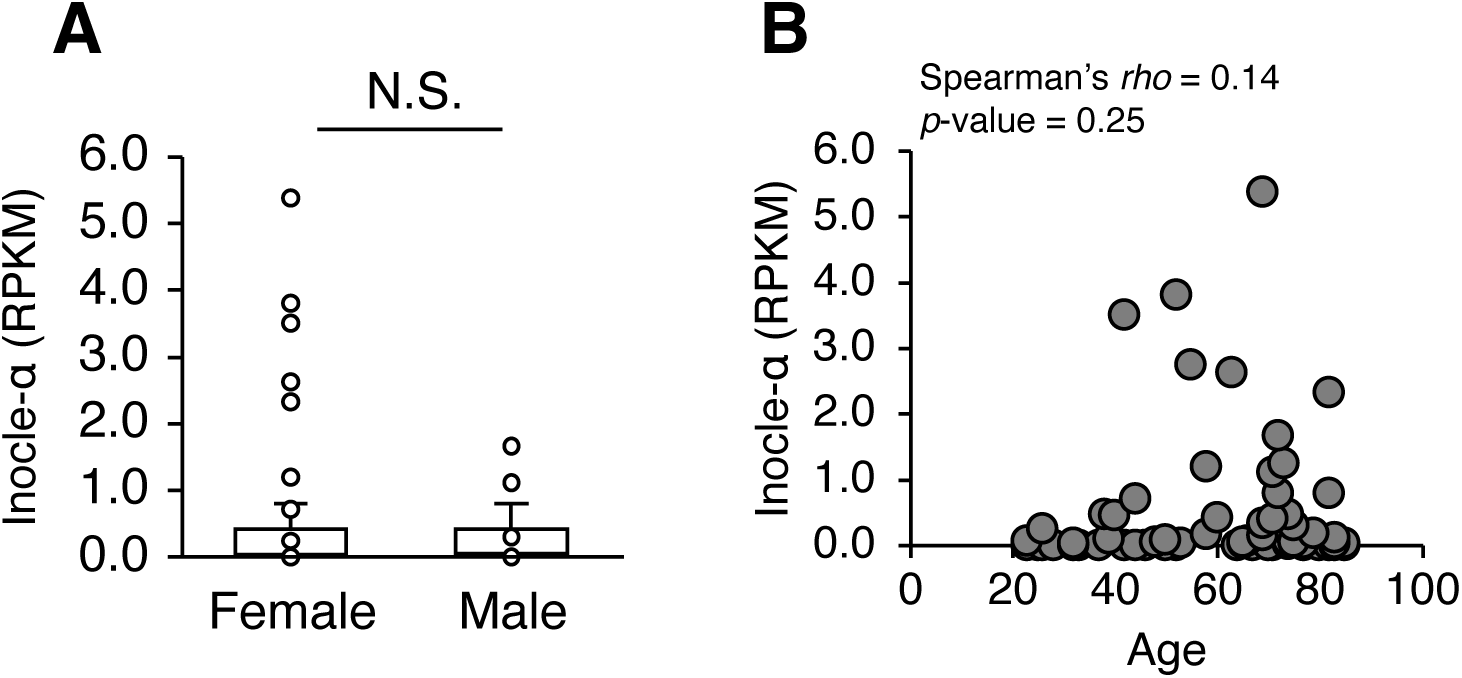
Associations of Inocles with sex and age. (A) Comparison of the abundance of Inocle-α in males and females. Box plots represent the inter-quartile range (IQR), and lines inside the box indicate the median. Whiskers show 1.5 IQR. N.S.= no significance based on the Wilcoxon rank-sum test. (B) Spearman’s *rho* between age and the abundance of Inocle-α.

**Figure S10.**
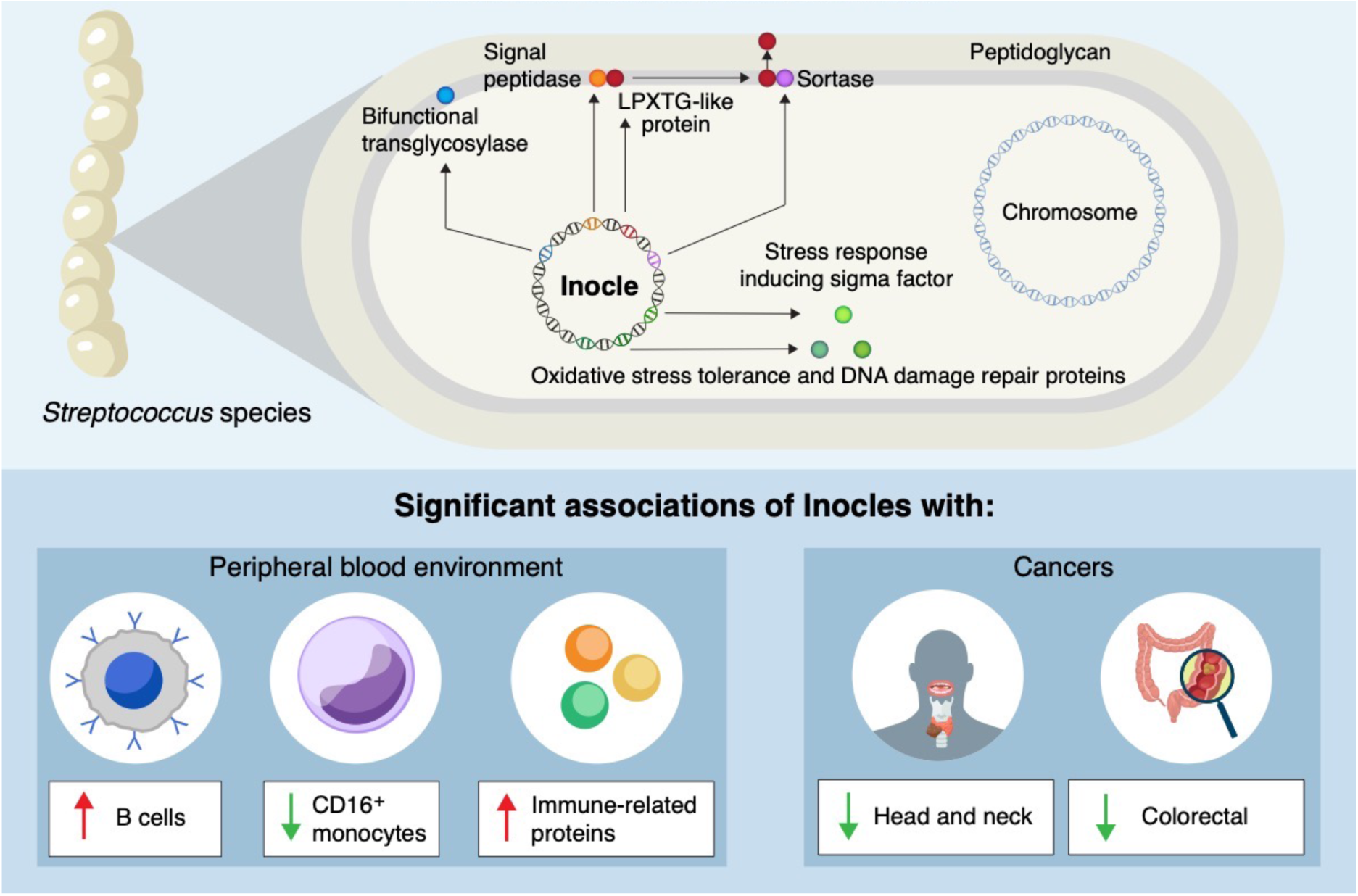
Functional characterization of Inocle. Schematic summary of functional annotation of Inocle based on gene annotation and significant association with human physiologies.

## METHODS

### Data availability

The short-read sequences, long-read sequences, and closed circular Inocles genomes are publicly available in DDBJ/GenBank/EMBL with accession number PRJDB18431. The sequences of InoC are available in zendo with DOI 10.5281/zenodo.13138926.

### Human saliva sample collection

This study was approved by the Ethics Committee of the University of Tokyo, National Cancer Center Hospital East, and Sam Ratulangi University. Informed consent was obtained from all the participants. Four saliva samples were collected from four Japanese participants to develop a preNuc method. For long-read metagenomics, salivary samples were collected from 46 Japanese, 9 Indonesian, and 1 Thailand participants. To estimate the prevalence of Inocles, we collected salivary samples from 68 Japanese and 20 Indonesian participants. In total, 45 salivary samples were collected from patients with HNC. Freshly collected salivary samples were immediately frozen at −80 °C and stored until use.

### DNA extraction from saliva samples

In the preNuc method, a 1 mL saliva sample was suspended in 500 μL of phosphate-buffered saline (PBS) and centrifuged at 12,000 × g for 10 min at 4 °C. The supernatant was discarded, and the pellet was suspended using 500 μL of Tris/MgCl buffer (40 mM Tris/HCl, 10 mM Mg/Cl2). Next, DNase I (50 units) and RNase I (5 μg/mL) (NIPPON GENE, Tokyo, Japan) were added and incubated for 30 min at 37 °C followed by centrifugation at 12,000 × g for 10 min at 4 °C. The supernatant was discarded; subsequently, the DNA was extracted and purified following a previously described method^45^.

For DNA extraction from small particle fractions in saliva, a 1 mL saliva sample was suspended in 1 ml of PBS centrifuged at 5,000 × g for 10 min at 4 °C. The supernatant was filtered using 0.45 μm PVDF pore membrane filters (Steriflip, Merck Millipore, Burlington, MA, USA). The filtrate was mixed with an equal volume of polyethylene glycol solution (20% PEG6000 in 2.5 M NaCl) and stored for 2 h at 4 °C. The solution was centrifuged at 20,000 × g for 45 min at 4 °C to collect small particles. After removing the supernatant, the pellets were suspended in 400 μL of TE20 buffer. The solution was processed following DNA extraction processes, a previously described method^45^.

### Long-read metagenomic sequencing and data processing

Library preparation was conducted using a Ligation Sequencing Kit (SQK-LSK114) following the protocol provided by the manufacturer (Oxford Nanopore Technologies, Oxford, UK). Sequencing was performed using PromethION (Oxford Nanopore Technologies) with R.10.4.1 flow cells. Base calling was performed using dorado-0.6.2, and long reads with an average Q score < 20 were removed using NanoFilt (v2.8.0)^46^. The quality-filtered long reads were mapped to the T2T genome^47^ using minimap2 (2.26-r1175)^48^ considering >90% identity and >85% alignment coverage and mapped reads were removed.

### Short-read metagenomic sequencing and data processing

The library was prepared using the TruSeq DNA Nano kit (Illumina, San Diego, CA, USA) or the TruSeq ChIP kit (Illumina), following the protocol provided by the manufacturers. Sequencing was performed using NovaSeq 6000 (Illumina, Inc., USA). Quality filtering of the NovaSeq reads was performed using fastp (v0.22.0)^49^ with the following options: n_base_limit 0 --trim_tail1 1 -- cut_mean_quality 20 --qualified_quality_phred 20 --unqualified_percent_limit 50 --length_required 50 --detect_adapter_for_pe 2 --trim_poly_g --cut_tail --cut_tail_window_size 1 –dedup. Quality-filtered reads were mapped to the T2T genome using bowtie2^50^ with default parameters, and mapped reads were removed.

### Isolation of the *Streptococcus* strain, including Inocle

To isolate the *Streptococcus* strains with Inocle_004, a fresh saliva sample (sample ID: m_5) was plated on De Man–Rogosa–Sharpe (MRS) agar plate (Becton, Dickinson and Company, USA) and incubated at 37 °C for one day. The presence of Inocle_004 was confirmed by Sanger sequencing of colony PCR product (226 bp) using Inocle_004 specific primer set (Inocle_004_forward 5’-AACGCCAGCTCTTCTGGATA-3’ and Inocle_004_reverse 5’-TGCTCGTGTAGGATCTGTCG-3’). The colony PCR was performed in 2 × KAPA HiFi DNA polymerase (Kapa Biosystems, USA) and 10μM primers under conditions of 3 min at 95 °C, 35 cycles of 98 °C for 20 sec, 60 °C for 30 sec, and 72 °C for 20 sec. For the whole genome shotgun sequencing of the bacterial strain with Inocle_004, the Inocle_004-positive strain was incubated in an MRS medium at 37 °C for one day, and genomic DNA was extracted using the DNeasy PowerSoil Pro Kit (QIAGEN, Germany) from a cultured medium. Then, the sequencing library was prepared using the NEBNext Ultra II DNA Library Prep Kit (New England Biolabs, USA). The genomic sequences were obtained using AVITI sequencer (Element Biosciences, San Diego, CA, USA).

The genomic assembly of the AVITI sequences was performed using subsampled reads (500,000 reads) with the default parameters of SPAdes^51^. The bacterial taxonomy of isolated strain was determined by GTDB-tk (v2.3.2)^52^. To calculate the Inocle-α/*S. salivarius* ratio in native saliva, metagenomic NovaSeq reads of the m_5 saliva sample (isolation source of Inocle_004) were mapped to the isolated *S. salivarius* and Inocle_004 genomes using bowtie2 (v2.4.1)^50^, and the RPKM was calculated from the RBP obtained using CoverM (v0.6.1) (--min-read-percent-identity 95 and --min-read-aligned-percent 85) normalized by the total number of reads. Subsequently, the Inocle-α/*S. salivarius* ratio was obtained by dividing the RPKM of Inocle-α by that of *S. salivarius*.

### Long-read metagenomic assembly

Long-read assembly was performed using Flye (2.9.2-b1786)^18^ with a --meta and --read-error 0.03. The circularity and sequence depth of each contig were obtained from the metaFlye output file. Initially, two Inocle contigs (Inocle_008 and Inocle_018) were assembled as linear contigs. To validate the circularity of these two contigs, a reference-guided assembly was performed. To purify the long reads of two Inocle genomes assembled as linear contigs (Inocle_008 and Inocle_018), long reads were mapped to initially assembled contigs considering >95% identity and >20% alignment coverage. Subsequently, mapped long reads were assembled using metaFlye, and circular contigs were obtained.

To verify the accuracy of long-read contigs, metagenomic short reads were obtained from two metagenomic DNA samples used to obtain the long reads. Short reads were assembled using MEGAHIT (v1.2.9)^53^. Short-read contigs were aligned to long-read contigs using minimap2 (2.26-r1175)^48^ with -cx asm20 parameter and obtained the similarity between short- and long-read contigs. All sequence statistics were obtained using Seqkit (v2.8.0)^54^.

### Classification of long-read contigs into microbial genetic elements

Small contigs <3kb and T2T genome-aligned (>80% identity and >20% alignment coverage) contigs were removed from the >10-depth long-read contigs. To identify the bacterial chromosomal contigs, rRNA marker gene sequences were predicted in long-read contigs using Barrnap (v0.9) (https://github.com/tseemann/barrnap) and Prokka (v 1.14.6)^23^. Other bacterial marker genes were predicted using fetchMG (v1.1)^55^ and contigs with bacterial marker genes were classified as bacterial chromosomal contigs. Binning analysis was performed using SemiBin2 (v2.0.2)^56^ with -- environment human_gut --sequencing-type=long_read parameters, and contigs in a Bin, including bacterial chromosomal contigs, were classified as potential chromosomal contigs. Potential phage contigs were predicted in non-bacterial chromosomal contigs using VirSorter2 (v2.2.4)^57^ with >0.9, and CheckV (v0.7.0)^58^ with contamination of <10%. Potential plasmid contigs from non-phage contigs were predicted using PlasClass (v0.1.1)^59^ with a >0.7 score. To identify eukaryotic viral and fungal contigs, non-plasmid contigs were aligned to RefSeq viral genomes (download date: 2023/07/06) and FugiDB (v64)^60^ using minimap2 (2.26-r1175)^48^ with parameter -cx asm20, >70% identity, and >50% alignment coverage. The remaining contigs were considered unclassified. The unclassified contigs were aligned to the plasmid^61^ and integrated phage genome database^4,5,8,9,13,62,63^.

ORFs of contigs were predicted using Prokka (v1.14.6)^23^ with default parameters. All ORFs were aligned to the UniRef90^64^ database using DIAMOND (v2.1.8)^65^ with --sensitive and an e-value<1e-10 options. Unclassified contigs with >50% ORFs unaligned with the UniRef90 database were classified as strong candidates for unrecognized genetic elements. Average amino acid identity (AAI)-based clustering was performed for contigs clustering^4^. All ORFs of the 85 strong candidate contigs of unrecognized genetic elements were subjected to “all vs. all aligned” by DIAMOND with --evalue 1e-5, --max-target-seqs 10000, --query-cover 50, and --subject-cover 50 options. All contig pairs satisfying a minimum of 80% shared ORFs, 30% average AAI, and > 10 shared ORFs were subjected to clustering analysis. Contig clustering was performed using MCL (v14-137) with -te 8 -I 2.0 –abc options. Contigs with >1.5-fold smaller or larger than the average contig size in the same cluster were treated as different clusters. Long-read mapping was performed using minimap2 (2.26-r1175)^48^ with map-ont option. The mapping results were visualized using the Integrative Genomics Viewer (IGV)^66^. To obtain the RPKM, first, we obtained the reads per base (RPB) using CoverM (v0.6.1) (https://github.com/wwood/CoverM) with --min-read-percent-identity 95 and --min-read-aligned-percent 85 options. Subsequently, the RBP was normalized to the total number of reads in each sample, and the RPKM was obtained.

### Genomic analysis of Inocles

Based on cumulative GC skew, the replication origin and terminus were predicted using iRep (v1.10)^67^. Genome visualization was performed using Proksee^68^. The insertion sequences were predicted using ISEscan (v1.7.2.3)^69^, considering the default parameters. All high-quality circular contigs having ORFs aligned to at least one InoC gene using DIAMOND with <1e-10 were identified as additional Inocle genomes. Average nucleotide identity was obtained using Pyani (v0.2.12)^70^ with the -m ANIb option. The PF of Inocles was obtained by ORF clustering using Roary (v3.13.0)^71^ with the -i 50 option. Based on the similarity of the PFs repertoire, a dendrogram was created using the ward.D2 method, considering the Euclidean distance calculated using the presence/absence information of the PFs. The short-read contigs assembled using MEGAHIT from the Inocle genome-reconstruction sample were aligned to Inocle genomes using minimap2 (2.26-r1175)^48^ with >99% identity and 500bp alignment length; fragmented positions in short-read contigs were determined based on aligned short-read contigs.

### Host bacterial prediction of Inocles

To identify the ORFs of Inocles homologous with host bacterial ORFs, all Inocle ORFs were aligned to the UniRef90^64^ by DIAMOND (v2.1.8)^65^ with an e-value <1e-10, and best-hit results were used for further analysis. The number of Inocle genes homologous to the host bacteria was calculated by summing the UniRef90 genera with the Inocle gene aligned.

Tetranucleotide frequency was obtained using CheckM (v1.1.3)^72^ from Inocles and chromosomes, plasmids, and phages of *Streptococcus*. All *Streptococcus* chromosomes, plasmids, and phages were obtained from the RefSeq database. The dendrogram based on the similarity of tetranucleotide frequency was obtained using the ward.D2 method based on the Manhattan distance.

Short reads of the bacterial pellet and 0.45 μm-filtered fraction from the saliva sample were mapped to the Inocle genome obtained from the same sample. The mapping was performed by bowtie2 (v2.4.1)^50^, and RPKM was calculated from the RBP obtained using CoverM (v0.6.1) (--min-read-percent-identity 95 and --min-read-aligned-percent 85) normalized to the total number of reads in each sample.

### Functional gene annotation of Inocles

InoC was identified via ORF clustering using cd-hit (v4.8.1)^73^ with -c 0.7 -n 5 -aS 0.5. The 3D structure of InoC was obtained using ColabFold (v1.5.5)^74^. The phylogenetic tree of InoC was constructed via multiple alignments using MAFFT (v7.505)^75^ following FastTree (v2.1.11)^76^. The phylogenetic tree of InoC was constructed using the iToL^77^. Functional annotation of ORFs was conducted using Prokka (v1.14.6)^23^. Mobilization-related genes were predicted using ICEscreen (v1.2.0)^78^ with default parameters. Subcellular localization of ORFs was predicted using Psort (v3.0)^79^ with the --positive option. The KEGG orthology annotations for the ORFs were performed using the eggNOG mapper (v2.1.11)^80^ (--sensmode ultra-sensitive), and the KEGG BRITE hierarchy was generated based on the assigned K numbers^81^. The Pfam domains and active sites of the signal peptidase in the ORFs were predicted using InterProScan^82^. Transmembrane regions in the cell wall/membrane-related ORFs were predicted using TMHMM-2.0^83^ with default parameters, and the protein 2D structure on the membrane was visualized using protter^84^. The LPXTG-like motif was predicted using CW-PRED^85^ with default parameters.

### Sequence analysis of deposited datasets

The deposited metagenomic sequences of HMP^19^ were downloaded from the NCBI SRA server. Low-quality reads were filtered using fastp^49^ with the following options --n_base_limit 0 --trim_tail1 1 --cut_mean_quality 20 --qualified_quality_phred 20 --unqualified_percent_limit 50 -- length_required 50 --detect_adapter_for_pe 2 --trim_poly_g --cut_tail --cut_tail_window_size 1 – dedup. The quality-filtered reads were mapped to the T2T genome using bowtie2^50^ with default parameters, and unmapped reads were extracted as filter-passed reads. Samples with >4 million filter-passed reads were used for further analysis. Filter-passed short reads were mapped to non-redundant InoC genes (clustered by 100% identity) using bowtie2 (v2.4.1)^50^, and the RPKM was calculated from the RBP obtained using CoverM (v0.6.1) (--min-read-percent-identity 95 and --min-read-aligned-percent 85) normalized by the total number of reads in each sample. The RPKM of each Inocle taxon was obtained by summing all RPKM of the InoC gene for each Inocle taxon. To confirm the transcripts of Inocle ORFs, metatranscriptome short reads were mapped to nucleotide sequences of Inocle PFs using bowtie2 (v2.4.1)^50^, and the RPKM was calculated from the RBP obtained using CoverM (v0.6.1) (--min-read-percent-identity 95 and --min-read-aligned-percent 85) normalized by the total number of reads in each sample.

To compare the abundance of Inocles, salivary metagenomic sequences from HCs and patients with PDAC^30^, RA^32^, or CRC^31^ in each cohort were collected from the NCBI SRA server; one sample was randomly selected from longitudinally sampled participants in the CRC study. Quality filtering and removal of human reads were performed using the same procedures as for the HMP study described above. Samples with >4 million filter-passed reads were used for comparison analysis.

### Single-cell RNA-seq analysis

A single-cell population dataset was selected based on a study by Kashima et al.^90^. Kashima et al. describe details of the experimental and bioinformatics processing^90^. In brief, the peripheral blood mononuclear cells (PBMCs) were subjected to library preparation for the scRNA-seq using 10× Genomics Chromium Next GEM Single Cell Multiome ATAC + Gene Expression according to the manufacturer’s user guide (CG000365 Rev A, CG000337 Rev A, 10× Genomics). The scRNA-seq was performed using NovaSeq 6000 (Illumina). Sequencing datasets were processed using Cell ranger ARC (Cellranger-arc-2.0.0, (default, intron-include mode, ref = GRCh38-2020-A-2.0.0)). Subsequently, NovaSeq reads were mapped to hg38 using STAR^86^ and obtained count data of human genes using Seurat (v4.1.1). Subsequently, count data was normalized and integrated to minimize batch effects using Harmony (v0.1.0). Cell clustering was performed using Seurat, and the clusters were manually annotated based on canonical markers described in the Azimuth reference^29^. GO enrichment was performed using ShinyGO 8.0^87^.

### Proteome analysis of human plasma samples

Blood samples and plasma fractions were collected for proteome analysis using Olink. Reactions with antibodies targeting 5,420 proteins were conducted following the protocol provided by the manufacturer (Olink Proteomics, Uppsala, Sweden) without bridging different experimental reactions for all samples. Sequence libraries were prepared following the protocol provided by the manufacturer (Olink Explore HT) followed by sequencing using a NovaSeq 6000 (Illumina). Sequence counts were transformed into normalized protein expression (NPX) values and further analyzed.

### Association analysis between Inocle-α, *S. salivarius*, blood cells, and plasma proteome

The abundance of Inocle-α was obtained as RPKM, and the abundance of *S. salivarius* was obtained using MetaPhlAn4 (v4.1.0)^88^. Correlation analysis was conducted using Spearman’s correlation coefficients with the psych package in RStudio (v2023.12.0+369).

## STATISTICAL ANALYSIS

### Statistical analysis

All statistical analyses were performed using RStudio (v2023.12.0+369). A two-tailed paired student’s t-test was used to compare the preNuc-treated and -untreated groups. Fisher’s exact test was used to compare the prevalence of Inocle-α and *S. salivarius* between the HNC and HC groups. For the association analysis between the Inocle-α and human diseases, the statistical test was performed using MaAsLin2^89^ with default parameters, except for the specific random effects. Data from HCs and patients with HNC were compared using MaAsLin2, with random effects of sex, age, smoking status, and cancer treatment status. Similarly, MaAsLin2, with random effects of sex and age, was performed to compare HCs and patients with PDAC, RA, and CRC in each cohort. Statistical results with *p* < 0.05 were considered significant.

